# GraphExperiment: infrastructure for high-dimensional quantitative data and associated networks in R/Bioconductor

**DOI:** 10.64898/2026.10.28.762566

**Authors:** Fabricio Almeida-Silva

**Affiliations:** Department of Plant Biotechnology and Bioinformatics, Ghent University, 9052 Ghent, Belgium; VIB Center for Plant Systems Biology, VIB, 9052 Ghent, Belgium

**Keywords:** graphs, systems biology, object-oriented programming, data integration

## Abstract

Networks are widely used to represent relationships among features (e.g. genes, proteins, and microbial taxa) and among observations (e.g. cells, samples, and species) in omics data, as well as their neighborhoods. The Bioconductor ecosystem provides users and developers with modular and interoperable software tools by relying on standard data structures that are reused across packages. However, Bioconductor lacks a standard data structure to store such networks together with the quantitative data from which they are inferred. Here, we present *GraphExperiment*, an R/Bioconductor package that provides an S4 class extending *SingleCellExperiment* with containers for feature-level (*rowGraphs*) and observation-level (*colGraphs*) networks stored as *igraph* objects. Networks are kept in sync with assays, row and column metadata, allowing the simultaneous manipulation of quantitative data and graphs. We demonstrate *GraphExperiment* in seven use cases covering bulk and single-cell transcriptomics, single-cell proteomics, microbiome, and comparative genomics data, with networks ranging from gene coexpression and protein co-abundance networks to cell-cell communication and phylogenetic networks. *GraphExperiment* is available on Bioconductor (https://bioconductor.org/packages/GraphExperiment).

## Introduction

Networks (or graphs) are ubiquitous in the analysis of omics data. At the feature level, researchers typically infer gene coexpression networks (Langfelder and Horvath 2008) and gene regulatory networks (Huynh-Thu et al. 2010) from transcriptomic data, protein co-abundance networks (Johnson et al. 2022) from proteomics data, and microbial co-occurrence networks (Kurtz et al. 2015) from microbiome data. At the observation level, standard applications of networks include nearest-neighbor graphs of cells for clustering of single-cell data (Xu and Su 2015), communication networks among cell types (Dimitrov et al. 2022), and species relationships with phylogenetic networks (Solís-Lemus and Ané 2016). In all these cases, the network is derived from (and/or interpreted with) a matrix of quantitative data and its associated metadata.

Much of the success of the Bioconductor project stems from shared data structures, ensuring seamless interoperability and a cohesive ecosystem of packages (Huber et al. 2015). For high-dimensional quantitative data, the `SummarizedExperiment` S4 class (Morgan et al. 2025) allows storing and accessing quantitative data with associated feature- and observation-level metadata in a single object. Since its release, the `SummarizedExperiment` class has been extended to include, among others, dimensionality reduction coordinates (the `SingleCellExperiment` class) (Amezquita et al. 2019), spatial coordinates and image data (the `SpatialExperiment` class) (Righelli et al. 2022), and phylogenetic trees (the `TreeSummarizedExperiment` class) (Huang et al. 2020). However, none of these classes stores networks. As a consequence, each package returns networks in its own format (e.g., edge lists, adjacency matrices, or package-specific lists), hindering interoperability and data integration. The `rowPairs` and `colPairs` slots of `SingleCellExperiment` objects store pairwise relationships as `SelfHits` objects, not as graphs, and `TreeSummarizedExperiment` objects store hierarchies (i.e., trees) only. Outside Bioconductor, `Seurat` (Hao et al. 2024) and `AnnData` (Virshup et al. 2024) objects store gene/cell graphs as sparse adjacency matrices rather than graph objects.

Here, we present *GraphExperiment*, an R/Bioconductor package that provides an S4 class to store high-dimensional quantitative data together with networks describing its features and observations. We describe the design of the class, and we demonstrate its use in seven use cases covering diverse omics data types and network types.

### Implementation

#### The GraphExperiment class

The `GraphExperiment` class extends `SingleCellExperiment` (Figure 1), hence inheriting all containers of the parent class, including *assays* (matrices of features by observations), *rowData* and *colData* (feature and observation metadata), *reduced-Dims* (low-dimensional representations), and *metadata*. It also inherits all methods, so packages that accept `SingleCellExperiment` or `SummarizedExperiment` objects accept `GraphExperiment` objects too. Relative to `SingleCellExperiment`, `GraphExperiment` adds two containers: *rowGraphs*, a list of graphs whose nodes are features, and *colGraphs*, a list of graphs whose nodes are observations.

**Figure 1.**
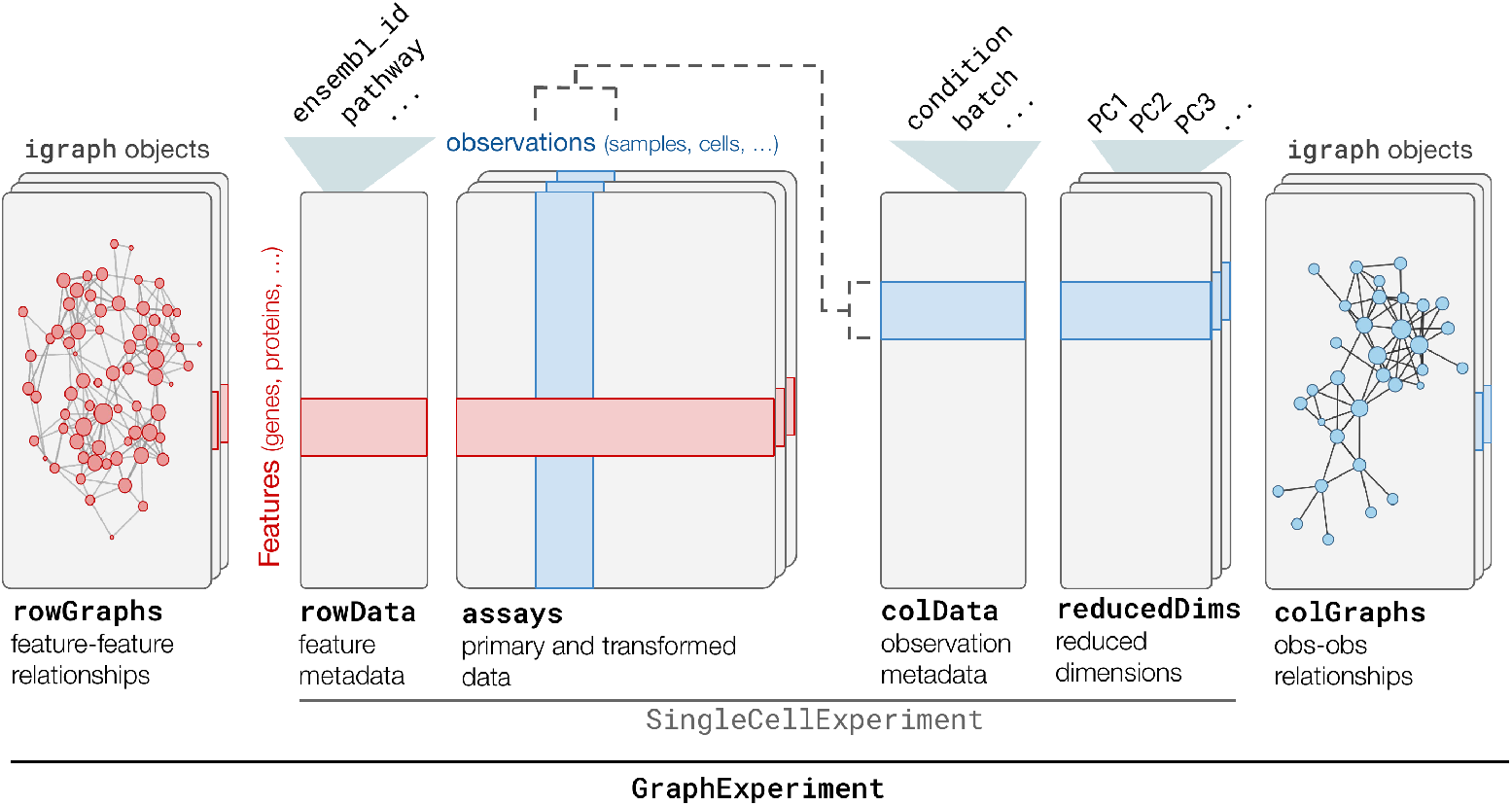
The GraphExperiment S4 class. Containers (or slots) inherited from `SingleCellExperiment` include *assays* (matrices of feature abundances across observations, e.g., genes x cells, proteins x samples, etc), *rowData* (metadata describing features), *colData* (metadata describing observations), and *reducedDims* (representations from dimensionality reduction algorithms, e.g., PCA and t-SNE). New containers include *rowGraphs* (graphs describing feature-feature relationships, such as gene-gene coexpression, protein-protein interactions, etc.) and *colGraphs* (graphs describing observation-observation relationships, such as cell-cell distances, species-species distances, etc.)

Graphs are stored as `igraph` objects (Csárdi and Nepusz 2006), the standard class for graph representation in R. Graphs can be directed or undirected, weighted or unweighted, and they can contain any number of node and edge attributes. We chose `igraph` objects over pair-wise relationships (as in the `rowPairs`/`colPairs` slots) for two reasons. First, `igraph` implements hundreds of graph algorithms (e.g., community detection, centrality measures, and edge- and graph-level statistics), and several packages already return `igraph` objects (e.g., *bluster* and *BioNERO* (A. Lun 2025; Almeida-Silva and Venancio 2021)). Second, `igraph` objects can be directly visualized and manipulated with popular packages such as *ggraph* and *tidygraph*.

### Construction, access, and validity

`GraphExperiment` objects can be created with the constructor function `GraphExperiment()`, which accepts the same arguments as `SingleCellExperiment()`, plus named lists of graphs in the arguments `rowGraphs` and `colGraphs`. Existing objects of class `SummarizedExperiment`, `RangedSummarizedExperiment`, and `SingleCellExperiment` (and, thus, of their subclasses, such as `TreeSummarizedExperiment`) can be coerced with the `as()` method. Graphs are accessed and replaced with `rowGraphs()` and `colGraphs()` (all graphs) or `rowGraph()` and `colGraph()` (a single graph, by name or index), in the same way as the `assays()` and `assay()` methods for `SummarizedExperiment` objects. Graph names can be accessed and replaced with the `rowGraphNames()` and `colGraphNames()` methods.

A validity method ensures that node names of each graph in *rowGraphs* and *colGraphs* match row names and column names, respectively. Hence, every feature (or observation) is a node, and every node is a feature (or observation). Features absent from an inferred network must be added as isolated nodes.

### Synchronization of graphs, metadata, and assays

`GraphExperiment` keeps graphs in sync with the rest of the object in two ways. First, it integrates node attributes with metadata. The `rowData()` and `colData()` methods return node attributes as additional columns named `<graph> <attribute>`, and the `rowGraph()` and `colGraph()` methods return graphs with *rowData* and *colData* variables as node attributes. For instance, module assignments of genes in a coexpression network named *gcn* are available in the column `gcn module` of *rowData*, and gene annotations in *rowData* can be used to color nodes when plotting the network. Second, subsetting rows or columns with the [ operator subsets *rowGraphs* or *colGraphs* to the induced subgraphs of the selected nodes, together with the assays and metadata.

## Results and Discussion

We demonstrate *GraphExperiment* in seven use cases (Table 1, Supplementary Data S1). To keep them short and reproducible, each use case uses a small example data set from an existing R/Bioconductor package or from a published study, and it runs in less than two minutes on a laptop (Intel i5-1345U processor; 16 GB RAM), with peak memory usage below 3 GB. The use cases focus on how networks are stored and used, not on new biological findings.

**Table 1.**
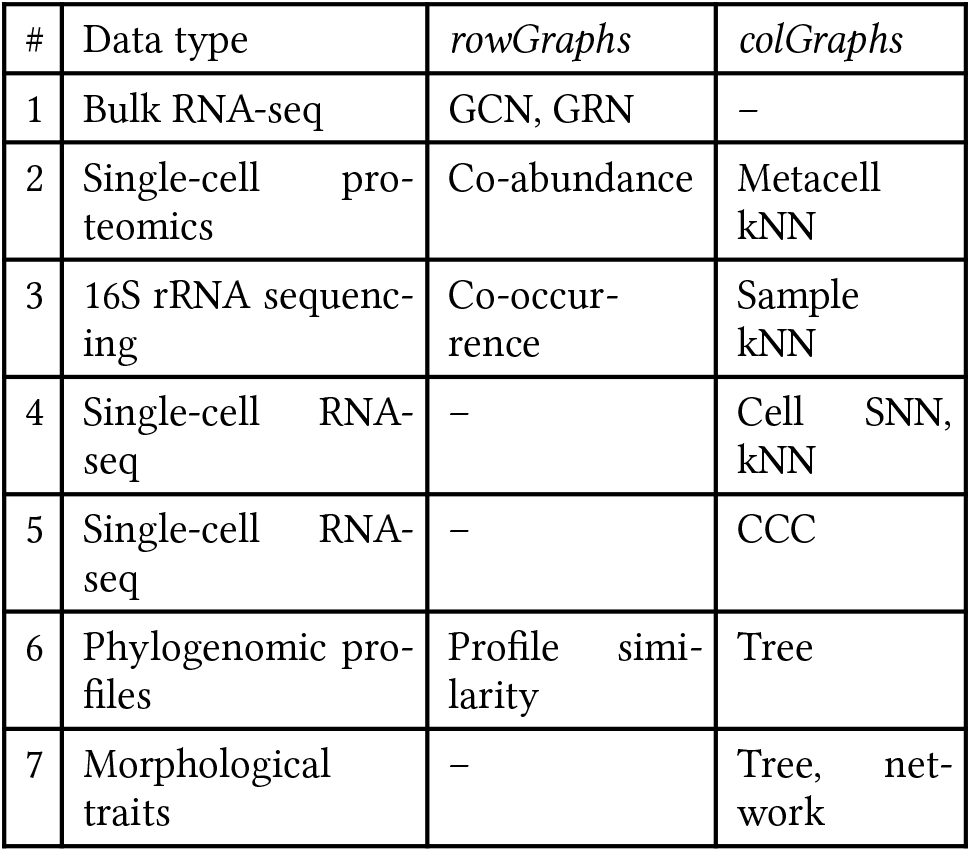
Overview of the use cases. GCN, gene coexpression network; GRN, gene regulatory network; kNN, k-nearest neighbor graph; SNN, shared nearest neighbor graph; CCC, cell-cell communication.

| # | Data type | *rowGraphs* | *colGraphs* |
|---|---|---|---|
| 1 | Bulk RNA-seq | GCN, GRN | – |
| 2 | Single-cell pro-teomics | Co-abundance | Metacell kNN |
| 3 | 16S rRNA sequenc-ing | Co-occur-rence | Sample kNN |
| 4 | Single-cell RNA-seq | – | Cell SNN, kNN |
| 5 | Single-cell RNA-seq | – | CCC |
| 6 | Phylogenomic pro-files | Profile simi-larity | Tree |
| 7 | Morphological traits | – | Tree, net-work |

In use case 1, we inferred a gene coexpression network from bulk RNA-seq data with the WGCNA algorithm (Langfelder and Horvath 2008) in *BioNERO* (Almeida-Silva and Venancio 2021), and stored the full network (i.e., a complete weighted graph) along with a gene regulatory network in *rowGraphs*. Subsetting the genes of one coexpression module subset both networks. We demonstrate how to store module assignments, hub status, and transcription factor annotations with the networks, and how to use standard `SummarizedExperiment` syntax to find transcription factors that are coexpression hubs and count their targets in the regulatory network. Use case 2 extends this approach to single-cell proteomics data (Khan et al. 2024; Vanderaa and Gatto 2023). We demonstrate how to aggregate cells into metacells to reduce sparsity deflating correlations (Morabito et al. 2023), and to infer networks representing protein co-abundances (*rowGraphs*) and built a kNN graph of distances between metacells (*colGraphs*).

Use case 3 integrates *GraphExperiment* with the miaverse ecosystem for microbiome data analysis (Borman et al. 2025). We coerced a `TreeSummarizedExperiment` object to a `GraphExperiment` object, and stored in *row-Graphs* taxon-taxon networks inferred with two methods for compositional data, namely SPIEC-EASI (Kurtz et al. 2015) and SparCC (Friedman and Alm 2012)). We also demonstrate how to compare both graphs, and how to infer and store a kNN graph of samples based on Aitchison distances in *colGraphs*.

Use cases 4 and 5 focus on single-cell RNA-seq data. In use case 4, we stored shared nearest-neighbor (SNN) and kNN graphs of cells, built with *scrapper* (A. T. L. Lun and Kancherla 2023) and *bluster* (A. Lun 2025), and added Louvain clusters as node attributes, which became available in *colData*. We then subset interneurons and re-clustered them using the induced subgraph to find subclusters. In use case 5, we inferred cell-cell communication with a consensus of methods in LIANA (Dimitrov et al. 2022), and stored the result as a directed graph of cell types (with ligand-receptor pairs as edge attributes) in a `GraphExperiment` object of pseudobulk profiles. This use case highlights that *colGraphs* can describe observations at any resolution, from cells to cell types.

Use cases 6 and 7 demonstrate how graphs can represent species relationships in comparative analyses. In use case 6, we use copy number variation of synteny clusters across eudicot plant species (Almeida-Silva et al. 2022) to demonstrate how to convert a phylogeny from a `phylo` object (Paradis and Schliep 2018) to an `igraph` object, and store a graph of species weighted by shared branch lengths in *colGraphs*. We then used this graph to test for phylogenetic signal in the copy number of each cluster, and stored a network of clusters with similar phylogenomic profiles in *rowGraphs*. Species relationships, however, are not always tree-like. Hybridization and introgression are better described by phylogenetic networks (Solís-Lemus and Ané 2016), and ignoring reticulation can bias comparative analyses (Morley, Lehmberg, and Kong 2026). Hence, in use case 7, we read phylogenetic networks of *Xiphophorus* fishes as DAGs, and show how to compute the expected covariance among species under Brownian motion (Bastide et al. 2018), and store covariance graphs derived from the tree and from the network in *colGraphs*. We then implement a function that fits phylogenetic generalized least squares models using any graph in *colGraphs*, and compare models fitted with each graph. This use case illustrates how developers can use *GraphExperiment* as the core data structure for new comparative methods based on graphs.

### Known limitations

Because node names must match row or column names, graphs cannot contain nodes that are not features or observations. In phylogenies and phylogenetic networks, for example, internal nodes represent ancestors, which are not observations. In use cases 6 and 7, we stored graphs of extant species in *colGraphs* and the full DAGs in the *metadata* slot. Future versions of *GraphExperiment* supporting extra nodes (not in assay rows or columns) are planned. Besides, `igraph` objects are stored in memory, and the number of edges in a full graph grows quadratically with the number of nodes. For large data sets, we recommend storing sparse graphs (e.g., after removing weak edges).

## Conclusion

*GraphExperiment* provides a standard data structure for quantitative data and associated networks in R/Bioconductor. By keeping networks in sync with assays and metadata, it simplifies network analyses for users and offers developers a common structure for the inputs and outputs of network inference and analysis methods. Our use cases show that the same class can store networks of genes, proteins, taxa, cells, cell types, and species. We expect *GraphExperiment* to become the standard container for networks in Bioconductor, as `SingleCellExperiment` became for single-cell data.

## Data availability

All data and code used in this article are available in its online Supplementary Material and at https://github.com/almeidasilvaf/GraphExperiment_paper.

