## Supplementary Data S1 for "GraphExperiment: infrastructure for high-dimensional quantitative data and associated networks in R/Bioconductor"

**Use cases for "GraphExperiment: infrastructure for high-dimensional quantitative data and associated networks in R/Bioconductor"**

Fabricio Almeida-Silva

2026

### Preface

This document is the Supplementary Data S1 of the article *GraphExperiment: infrastructure for high-dimensional quantitative data and associated networks in R/Bioconductor*. It contains the use cases described in the article, with all code needed to reproduce them. Code and data are also available at https://github.com/almeidasilvaf/GraphExperiment_paper.

**GraphExperiment** (Almeida-Silva 2026) extends the `SingleCellExperiment` class with two new slots: `rowGraphs`, which stores graphs describing how features (e.g., genes, proteins, taxa) relate to each other, and `colGraphs`, which stores graphs describing how observations (e.g., samples, cells, cell types, species) relate to each other. Here, we demonstrate how **GraphExperiment** can be used with different omics data types through a series of short use cases, written in the style of package vignettes. All use cases use small example data sets distributed with other R/Bioconductor packages, and they can be run on a laptop.

Table 1: Use cases in this document.

| Use case | Data type | Graphs | Slot |
|---|---|---|---|
| 1 | Bulk RNA-seq (maize tissues) | Gene coexpression and regulatory networks | `rowGraphs` |
| 2 | Single-cell proteomics (EMT time course) | Protein co-abundance network; metacell kNN graph | `rowGraphs`, `colGraphs` |
| 3 | 16S rRNA amplicon sequencing (human skin) | Taxon association networks (SPIEC-EASI, SparCC); sample kNN graph | `rowGraphs`, `colGraphs` |
| 4 | scRNA-seq (mouse brain) | Cell-cell SNN and kNN graphs | `colGraphs` |
| 5 | scRNA-seq (human pancreas) | Cell type communication network | `colGraphs` |
| 6 | Comparative genomics (eudicots) | Phylogenetic profile similarity network; species graph from a phylogeny | `rowGraphs`, `colGraphs` |
| 7 | Comparative phenotypic data (*Xiphophorus*) | Species graphs from a tree and from a phylogenetic network | `colGraphs` |

#### Installation

To reproduce the analyses in this book, you will need to install the following packages:

```r
install.packages("BiocManager")
pkgs <- c(
    "remotes", "tidyverse", "ggraph", "patchwork", "here", "nipals",
    "GraphExperiment", "BioNERO", "scpdata", "QFeatures", "org.Hs.eg.db",
    "limma", "mia", "scater", "scuttle", "scrapper", "bluster", "scRNAseq",
    "syntenet", "ComplexHeatmap", "zdk123/SpiecEasi", "saezlab/liana"
)

BiocManager::install(pkgs)
```

### Use case 1: gene coexpression and regulatory networks

Here, we will use **GraphExperiment** to store gene-gene networks inferred from bulk RNA-seq data. We will use the maize (*Zea mays*) gene expression data set distributed with the **BioNERO** package (Almeida-Silva and Venancio 2021), which contains 28 samples from different tissues (Shin et al. 2020). From these data, we will infer a gene coexpression network (GCN) and a gene regulatory network (GRN), and store both in the `rowGraphs` slot of a `GraphExperiment` object.

```
library(here)
library(GraphExperiment)
library(BioNERO)
library(SummarizedExperiment)
library(tidyverse)
library(ggraph)
library(patchwork)

set.seed(123) # for reproducibility
```

#### Data description and processing

The expression data are stored in a `SummarizedExperiment` object with a gene expression matrix (in TPM) and sample metadata. **BioNERO** also contains a data frame with maize transcription factors (TFs), which we will use to infer the GRN.

```
# Load data
data(zma.se)
data(zma.tfs)

zma.se
```

```
class: SummarizedExperiment
dim: 10802 28
metadata(0):
assays(1): ''
rownames(10802): ZeamMp030 ZeamMp044 ... Zm00001d054106 Zm00001d054107
rowData names(0):
colnames(28): SRX339756 SRX339757 ... SRX2792103 SRX2792104
colData names(1): Tissue
```

```
table(zma.se$Tissue)
```

```

    endosperm        pollen whole_seedling
            8             5             15
```

```
head(zma.tfs)
```

```
             Gene Family
6  Zm00001d022525    Dof
25 Zm00001d037605   GATA
28 Zm00001d049540    NAC
45 Zm00001d042287    MYB
46 Zm00001d042288    NAC
54 Zm00001d039371    TCP
```

To keep this example lightweight, we will remove lowly expressed genes and keep only the top 2,000 most variable genes. We will also use `exp_preprocess()` to remove outlier samples and adjust for confounders.

```
# Preprocess data and keep only the top 2,000 most variable genes
exp <- exp_preprocess(
    zma.se, min_exp = 10, variance_filter = TRUE, n = 2000
)

exp
```

```
class: SummarizedExperiment
dim: 2000 27
metadata(0):
assays(1): ''
rownames(2000): ZeamMp030 ZeamMp092 ... Zm00001d054093 Zm00001d054107
rowData names(0):
colnames(27): SRX339756 SRX339757 ... SRX2792103 SRX2792104
colData names(1): Tissue
```

#### Gene coexpression network inference

Next, we will infer a GCN with the WGCNA algorithm (Langfelder and Horvath 2008) as implemented in **BioNERO**. First, we need to pick the power $\beta$ to which correlation coefficients will be raised to make the network fit a scale-free topology.

```
# Pick power for scale-free topology fit
sft <- SFT_fit(exp, net_type = "signed hybrid", cor_method = "pearson")
```

```
   Power SFT.R.sq    slope truncated.R.sq mean.k. median.k. max.k.
1      3 0.293000  0.27100         0.1180   384.0     386.0    689
2      4 0.000141 -0.00465        -0.2750   290.0     272.0    584
3      5 0.210000 -0.20100         0.0542   227.0     202.0    509
4      6 0.427000 -0.35900         0.2990   184.0     155.0    452
5      7 0.583000 -0.48400         0.4780   153.0     121.0    407
6      8 0.665000 -0.58300         0.5720   129.0      96.0    370
7      9 0.697000 -0.66500         0.6110   111.0      77.8    339
8     10 0.786000 -0.71800         0.7260    95.8      64.1    313
9     11 0.787000 -0.77600         0.7310    83.8      53.4    290
10    12 0.821000 -0.82800         0.7810    73.9      44.7    270
11    13 0.857000 -0.86700         0.8290    65.6      37.5    252
12    14 0.884000 -0.89500         0.8660    58.6      31.5    236
13    15 0.890000 -0.91400         0.8710    52.7      26.7    221
14    16 0.884000 -0.93900         0.8630    47.6      22.9    208
15    17 0.886000 -0.96300         0.8630    43.1      19.7    196
16    18 0.896000 -0.97500         0.8740    39.2      17.0    185
17    19 0.905000 -0.98400         0.8840    35.8      14.8    175
18    20 0.914000 -0.99300         0.8930    32.8      12.8    166
```

```r
sft$power
```

```
[1] 12
```

Now, we can infer the GCN.

```r
# Infer the GCN
gcn <- exp2gcn(
    exp, net_type = "signed hybrid", SFTpower = sft$power,
    cor_method = "pearson"
)
```

```
..connectivity..
..matrix multiplication (system BLAS)..
..normalization..
..done.
```

```r
# Number of genes per module
table(gcn$genes_and_modules$Modules)
```

```

    black      cyan darkgreen   darkred      grey    purple       red
      518       878       245       222        56        42        39
```

The object returned by `exp2gcn()` contains the adjacency matrix of the full network, which we will convert to an `igraph` object. Note that we will **not** filter edges here. The adjacency matrix represents a complete, weighted graph, with edge weights representing the (soft-thresholded) coexpression between gene pairs. Then, we will add module assignments and intramodular connectivities as node attributes.

```r
# Convert adjacency matrix to an igraph object
gcn_graph <- graph_from_adjacency_matrix(
    gcn$adjacency_matrix, mode = "undirected", weighted = TRUE, diag = FALSE
)

# Add module assignments and intramodular degree as node attributes
nodes <- V(gcn_graph)$name
modules <- setNames(gcn$genes_and_modules$Modules, gcn$genes_and_modules$Genes)
V(gcn_graph)$module <- modules[nodes]
V(gcn_graph)$kWithin <- gcn$kIN[nodes, "kWithin"]

# Identify hub genes and add them as node attributes
hubs <- get_hubs_gcn(exp, gcn)
V(gcn_graph)$is_hub <- nodes %in% hubs$Gene

gcn_graph
```

```
IGRAPH a45d9dc UNW- 2000 1621533 --
+ attr: name (v/c), module (v/c), kWithin (v/n), is_hub (v/l), weight
| (e/n)
+ edges from a45d9dc (vertex names):
 [1] ZeamMp030--ZeamMp092 ZeamMp030--ZeamMp116 ZeamMp030--ZemaCp003
 [4] ZeamMp030--ZemaCp004 ZeamMp030--ZemaCp005 ZeamMp030--ZemaCp008
 [7] ZeamMp030--ZemaCp009 ZeamMp030--ZemaCp021 ZeamMp030--ZemaCp022
[10] ZeamMp030--ZemaCp024 ZeamMp030--ZemaCp026 ZeamMp030--ZemaCp027
[13] ZeamMp030--ZemaCp028 ZeamMp030--ZemaCp029 ZeamMp030--ZemaCp034
```

```
[16] ZeamMp030--ZemaCp035 ZeamMp030--ZemaCp036 ZeamMp030--ZemaCp040
[19] ZeamMp030--ZemaCp044 ZeamMp030--ZemaCp045 ZeamMp030--ZemaCp051
+ ... omitted several edges
```

The result is an `igraph` object with vertex (or node) attributes `name`, `module`, `kWithin`, and `is_hub`, and an edge attribute `weight`.

#### Gene regulatory network inference

Now, we will infer a GRN using the 'wisdom of the crowds' approach implemented in `exp2grn()`, which combines the results of GENIE3, ARACNE, and CLR (Huynh-Thu et al. 2010; Margolin et al. 2006; Faith et al. 2007) and selects the top edges that make the network best fit a scale-free topology.

```
# Get TFs among the top 2,000 most variable genes
tfs <- intersect(zma.tfs$Gene, rownames(exp))
length(tfs)
```

```
[1] 31
```

```
# Infer GRN
grn_edges <- exp2grn(exp, regulators = tfs, nTrees = 1000)
head(grn_edges)
```

```
          Regulator          Target
285 Zm00001d041474 Zm00001d012007
280 Zm00001d041474 Zm00001d006602
291 Zm00001d041474 Zm00001d027374
282 Zm00001d041474 Zm00001d008691
288 Zm00001d041474 Zm00001d015811
304 Zm00001d042263 Zm00001d045042
```

GRNs typically include only a subset of the genes, as genes that are not regulated by any TF (or that are not TFs) are not included in the edge list. However, `GraphExperiment` requires that nodes in row graphs match row names *exactly*. Thus, when creating the `igraph` object, we will include all genes as nodes, and genes that are not in the GRN will be represented as isolated nodes.

```
# Create a directed graph from the edge list, including all genes as nodes
grn_graph <- graph_from_data_frame(
    grn_edges, directed = TRUE, vertices = data.frame(name = rownames(exp))
)

grn_graph
```

```
IGRAPH cd5948b DN-- 2000 247 --
+ attr: name (v/c)
+ edges from cd5948b (vertex names):
 [1] Zm00001d041474->Zm00001d012007 Zm00001d041474->Zm00001d006602
 [3] Zm00001d041474->Zm00001d027374 Zm00001d041474->Zm00001d008691
 [5] Zm00001d041474->Zm00001d015811 Zm00001d042263->Zm00001d045042
 [7] Zm00001d041474->Zm00001d018986 Zm00001d041474->Zm00001d014773
 [9] Zm00001d041474->Zm00001d011541 Zm00001d041474->Zm00001d006942
[11] Zm00001d054038->Zm00001d032547 Zm00001d041474->Zm00001d018668
[13] Zm00001d025339->Zm00001d035115 Zm00001d041474->Zm00001d015279
[15] Zm00001d042263->Zm00001d043382 Zm00001d044315->Zm00001d043497
+ ... omitted several edges
```

#### Storing networks in a GraphExperiment object

Now that we have both networks, we can store them in a GraphExperiment object along with the expression data and sample metadata. We will also add gene-level metadata indicating whether each gene encodes a TF.

```
# Add gene-level metadata to `rowData`
rowData(exp)$is_TF <- rownames(exp) %in% zma.tfs$Gene
rowData(exp)$TF_family <- zma.tfs$Family[match(rownames(exp), zma.tfs$Gene)]

# Create a `GraphExperiment` object
ge <- GraphExperiment(
    assays = assays(exp),
    colData = colData(exp),
    rowData = rowData(exp),
    rowGraphs = list(gcn = gcn_graph, grn = grn_graph)
)

ge
```

```
class: GraphExperiment
dim: 2000 27
metadata(0):
assays(1): ''
rownames(2000): ZeamMp030 ZeamMp092 ... Zm00001d054093 Zm00001d054107
rowData names(5): is_TF TF_family gcn__module gcn__kWithin gcn__is_hub
colnames(27): SRX339756 SRX339757 ... SRX2792103 SRX2792104
colData names(1): Tissue
reducedDimNames(0):
mainExpName: NULL
altExpNames(0):
rowGraphs(2): gcn grn
colGraphs(0):
```

```
rowGraphNames(ge)
```

```
[1] "gcn" "grn"
```

Note that node attributes in each graph are automatically added to rowData, with column names in the format <graph name>__<attribute>. This way, module assignments, intramodular degree, and hub status can be accessed (and used, e.g., for subsetting) as any other feature-level metadata.

```
rowData(ge)
```

```
DataFrame with 2000 rows and 5 columns
                   is_TF   TF_family gcn__module gcn__kWithin gcn__is_hub
               <logical> <character> <character>    <numeric>   <logical>
ZeamMp030          FALSE          NA       black    27.178194       FALSE
ZeamMp092          FALSE          NA      purple     1.567267       FALSE
ZeamMp108          FALSE          NA      purple     0.214523       FALSE
ZeamMp116          FALSE          NA      purple     4.144444       FALSE
ZeamMp158          FALSE          NA        grey     0.175042       FALSE
...                  ...         ...         ...          ...         ...
Zm00001d054065     FALSE          NA        cyan     201.6571       FALSE
Zm00001d054084     FALSE          NA        cyan     222.0207       FALSE
```

```
Zm00001d054089     FALSE          NA        cyan     175.8362       FALSE
Zm00001d054093     FALSE          NA       black      18.8067       FALSE
Zm00001d054107     FALSE          NA        cyan      92.0203       FALSE
```

The opposite also holds: when graphs are extracted with `rowGraph()`, variables in `rowData` are added as node attributes. In other words, features and their metadata are always synchronized, regardless of whether one is working with the assay or with the graphs.

```
# Extract GRN with `rowData` variables as node attributes
grn_graph <- rowGraph(ge, "grn")
grn_graph
```

```
IGRAPH cd5948b DN-- 2000 247 --
+ attr: name (v/c), is_TF (v/l), TF_family (v/c)
+ edges from cd5948b (vertex names):
 [1] Zm00001d041474->Zm00001d012007 Zm00001d041474->Zm00001d006602
 [3] Zm00001d041474->Zm00001d027374 Zm00001d041474->Zm00001d008691
 [5] Zm00001d041474->Zm00001d015811 Zm00001d042263->Zm00001d045042
 [7] Zm00001d041474->Zm00001d018986 Zm00001d041474->Zm00001d014773
 [9] Zm00001d041474->Zm00001d011541 Zm00001d041474->Zm00001d006942
[11] Zm00001d054038->Zm00001d032547 Zm00001d041474->Zm00001d018668
[13] Zm00001d025339->Zm00001d035115 Zm00001d041474->Zm00001d015279
[15] Zm00001d042263->Zm00001d043382 Zm00001d044315->Zm00001d043497
+ ... omitted several edges
```

#### Exploring the networks

Since all data are stored in a single object, subsetting is straightforward. For example, let's subset the object to keep only genes in the module with the largest number of TFs. Subsetting a `GraphExperiment` object subsets assays, `rowData`, and all graphs in `rowGraphs` simultaneously.

```
# Get the module with the largest number of TFs
tfs_per_module <- table(rowData(ge)$gcn__module[rowData(ge)$is_TF])
tfs_per_module
```

```
    black      cyan darkgreen   darkred      grey       red 
       14         9         2         2         1         3 
```

```
top_module <- names(which.max(tfs_per_module))

# Subset the object to keep only genes in this module
ge_mod <- ge[rowData(ge)$gcn__module == top_module, ]
ge_mod
```

```
class: GraphExperiment
dim: 518 27
metadata(0):
assays(1): ''
rownames(518): ZeamMp030 Zm00001d000019 ... Zm00001d053845
  Zm00001d054093
rowData names(5): is_TF TF_family gcn__module gcn__kWithin gcn__is_hub
colnames(27): SRX339756 SRX339757 ... SRX2792103 SRX2792104
colData names(1): Tissue
reducedDimNames(0):
```

```
mainExpName: NULL
altExpNames(0):
rowGraphs(2): gcn grn
colGraphs(0):
```

Note that both graphs were filtered to keep only genes in the module.

```
# Check number of nodes per graph
lapply(rowGraphs(ge_mod), vcount)
```

```
$gcn
[1] 518

$grn
[1] 518
```

Finally, let's visualize the coexpression and regulatory networks for genes in this module, highlighting TFs and hub genes. For visualization purposes, we will only display the strongest coexpression edges.

```
# Extract graphs from the subset object (with `rowData` as node attributes)
g_gcn <- rowGraph(ge_mod, "gcn")
g_grn <- rowGraph(ge_mod, "grn")

# Keep only top 5% edges in the GCN (for visualization only)
w <- E(g_gcn)$weight
g_gcn <- delete_edges(g_gcn, which(w < quantile(w, 0.95)))

# Remove isolated nodes from both graphs (for visualization only)
g_gcn <- delete_vertices(g_gcn, which(degree(g_gcn) == 0))
g_grn <- delete_vertices(g_grn, which(degree(g_grn) == 0))

# Plot GCN, with node size proportional to intramodular degree
p_gcn <- ggraph(g_gcn, layout = "fr") +
    geom_edge_link(aes(alpha = weight), color = "grey60", show.legend = FALSE) +
    geom_node_point(aes(size = kWithin, color = is_hub)) +
    scale_color_manual(values = c(`FALSE` = "grey40", `TRUE` = "firebrick")) +
    theme_graph(base_family = "sans") +
    labs(
        title = paste0("Coexpression network (module '", top_module, "')"),
        color = "Hub gene", size = "Intramodular degree"
    )

# Plot GRN, highlighting TFs
p_grn <- ggraph(g_grn, layout = "fr") +
    geom_edge_link(
        arrow = arrow(length = unit(1.5, "mm")), end_cap = circle(1.5, "mm"),
        color = "grey60"
    ) +
    geom_node_point(aes(color = is_TF, size = is_TF)) +
    scale_color_manual(values = c(`FALSE` = "grey40", `TRUE` = "steelblue")) +
    scale_size_manual(values = c(`FALSE` = 1.5, `TRUE` = 4)) +
    theme_graph(base_family = "sans") +
    labs(
        title = paste0("Regulatory network (module '", top_module, "')"),
        color = "TF", size = "TF"
    )

wrap_plots(p_gcn, p_grn, nrow = 1) +
```

```r
    plot_annotation(tag_levels = "A") &
    theme(plot.title = element_text(size = 13))
```

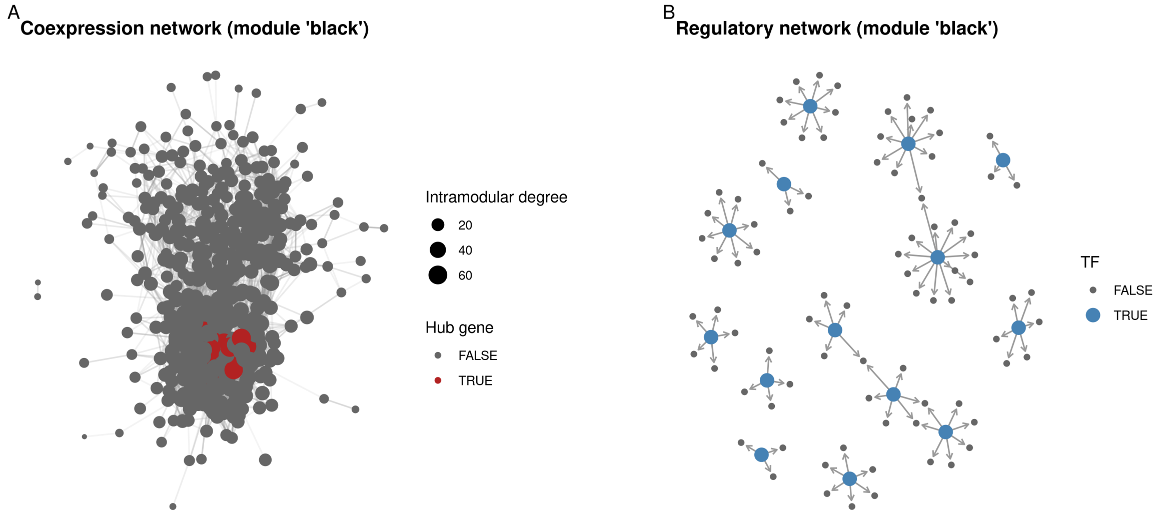

Because module assignments, hub status, and TF annotation travel together with the graphs, any downstream analysis (e.g., finding TFs that are hubs in both networks) can be done with standard `SummarizedExperiment` syntax.

```r
# TFs that are also GCN hubs, with their out-degree in the GRN
rowData(ge) |>
    as.data.frame() |>
    rownames_to_column("gene") |>
    filter(is_TF, gcn__is_hub) |>
    mutate(grn_outdegree = degree(rowGraph(ge, "grn"), gene, mode = "out")) |>
    select(gene, TF_family, gcn__module, gcn__kWithin, grn_outdegree)
```

```
              gene TF_family gcn__module gcn__kWithin grn_outdegree
1 Zm00001d025339 S1Fa-like         red     10.37434            12
2 Zm00001d041474       LSD     darkred     25.68034            16
3 Zm00001d042263      bHLH         red     10.78855             7
```

#### Session info

This document was created under the following conditions:

```
- Session info ---------------------------------------------------------------
 setting  value
 version  R version 4.5.2 (2025-10-31)
 os       Ubuntu 22.04.5 LTS
 system   x86_64, linux-gnu
 ui       X11
 language (EN)
 collate  en_US.UTF-8
 ctype    en_US.UTF-8
 tz       Europe/Brussels
 date     2026-10-24
 pandoc   3.6.3 @ /opt/quarto/bin/tools/ (via rmarkdown)
```

```
 quarto   1.8.25 @ /usr/local/bin/quarto

- Packages -------------------------------------------------------------------
 package              * version date (UTC) lib source
 Biobase              * 2.70.0  2025-10-29 [1] Bioconductor 3.22 (R 4.5.2)
 BiocGenerics         * 0.56.0  2025-10-29 [1] Bioconductor 3.22 (R 4.5.2)
 BioNERO              * 1.18.0  2025-10-29 [1] Bioconductor 3.22 (R 4.5.2)
 dplyr                * 1.1.4   2023-11-17 [1] CRAN (R 4.5.2)
 forcats              * 1.0.1   2025-09-25 [1] CRAN (R 4.5.2)
 generics             * 0.1.4   2025-05-09 [1] CRAN (R 4.5.2)
 GenomicRanges        * 1.62.1  2025-12-08 [1] Bioconductor 3.22 (R 4.5.2)
 ggplot2              * 4.0.1   2025-11-14 [1] CRAN (R 4.5.2)
 ggraph               * 2.2.2   2025-08-24 [1] CRAN (R 4.5.2)
 GraphExperiment      * 1.1.2   2026-09-08 [1] Bioconductor
 here                 * 1.0.2   2025-09-15 [1] CRAN (R 4.5.2)
 igraph               * 2.2.1   2025-10-27 [1] CRAN (R 4.5.2)
 IRanges              * 2.44.0  2025-10-29 [1] Bioconductor 3.22 (R 4.5.2)
 lubridate            * 1.9.4   2024-12-08 [1] CRAN (R 4.5.2)
 MatrixGenerics       * 1.22.0  2025-10-29 [1] Bioconductor 3.22 (R 4.5.2)
 matrixStats          * 1.5.0   2025-01-07 [1] CRAN (R 4.5.2)
 patchwork            * 1.3.2   2025-08-25 [1] CRAN (R 4.5.2)
 purrr                * 1.2.1   2026-01-09 [1] CRAN (R 4.5.2)
 readr                * 2.1.6   2025-11-14 [1] CRAN (R 4.5.2)
 S4Vectors            * 0.48.0  2025-10-29 [1] Bioconductor 3.22 (R 4.5.2)
 Seqinfo              * 1.0.0   2025-10-29 [1] Bioconductor 3.22 (R 4.5.2)
 SingleCellExperiment * 1.32.0  2025-10-29 [1] Bioconductor 3.22 (R 4.5.2)
 stringr              * 1.6.0   2025-11-04 [1] CRAN (R 4.5.2)
 SummarizedExperiment * 1.40.0  2025-10-29 [1] Bioconductor 3.22 (R 4.5.2)
 tibble               * 3.3.1   2026-01-11 [1] CRAN (R 4.5.2)
 tidyr                * 1.3.2   2025-12-19 [1] CRAN (R 4.5.2)
 tidyverse            * 2.0.0   2023-02-22 [1] CRAN (R 4.5.2)

 [1] /home/faalm/R/x86_64-pc-linux-gnu-library/4.5
 [2] /usr/local/lib/R/site-library
 [3] /usr/lib/R/site-library
 [4] /usr/lib/R/library
 * -- Packages attached to the search path.

------------------------------------------------------------------------------
```

### Use case 2: protein co-abundance networks from single-cell proteomics

Here, we will use **GraphExperiment** to store a protein co-abundance network inferred from mass spectrometry-based single-cell proteomics (SCP) data. Co-abundance networks are typically inferred with the WGCNA algorithm (Langfelder and Horvath 2008), which has been widely used in large-scale proteomics studies (Johnson et al. 2022; Seyfried et al. 2017), because soft-thresholding and topological overlap-based module detection are more robust than simple correlation thresholds.

We will use the data set from Khan et al. (2024), available in the **scpdata** package (Vanderaa and Gatto 2023a), which contains SCP data for 421 MCF-10A cells undergoing epithelial-mesenchymal transition (EMT) triggered by TGF-$\beta$, with cells collected at days 0, 3, and 9.

SCP data (as scRNA-seq data) are very sparse, which deflates correlations and makes co-abundance network inference unreliable. A common approach to handle sparsity is to aggregate similar cells into *metacells* before network inference (Morabito et al. 2023), which is what we will do here.

```
library(here)
library(GraphExperiment)
library(scpdata)
library(QFeatures)
library(BioNERO)
library(bluster)
library(tidyverse)
library(ggraph)

set.seed(123) # for reproducibility
```

#### Data description

The data are stored in a `QFeatures` object (Gatto and Vanderaa 2025) with quantitative data at the PSM, peptide, and protein levels. Here, we will only use the protein-level data, without imputation of missing values.

```
# Load data
qf <- khan2023()
```

```
qf
```

```
An instance of class QFeatures (type: scp) with 47 sets:

 [1] eSK233: SingleCellExperiment with 5951 rows and 16 columns
 [2] eSK234: SingleCellExperiment with 6375 rows and 16 columns
 [3] eSK235: SingleCellExperiment with 6261 rows and 16 columns
 ...
 [45] peptides: SingleCellExperiment with 10055 rows and 421 columns
 [46] proteins_imputed: SingleCellExperiment with 4096 rows and 421 columns
 [47] proteins_unimputed: SingleCellExperiment with 4096 rows and 421 columns
```

```r
# Get protein-level data with sample metadata
prot <- getWithColData(qf, "proteins_unimputed")
rm(qf) # free memory

prot
```

```
class: SingleCellExperiment
dim: 4096 421
metadata(0):
assays(1): ''
rownames(4096): A0A075B6I4 A0A0A0MS14 ... Q9Y6X9-2 W5XKT8-3
rowData names(1): protein
colnames(421): eSK234RI10 eSK239RI5 ... eSK238RI15 eSK238RI16
colData names(7): Set Channel ... sortday digest
reducedDimNames(0):
mainExpName: NULL
altExpNames(0):
```

Let's have a look at the available *colData* variables, and the proportion of missing values in our *assay*.

```r
# Quick check on colData variables
colData(prot)
```

```
DataFrame with 421 rows and 7 columns
                   Set     Channel SampleAnnotation  SampleType     lcbatch
           <character> <character>      <character> <character> <character>
eSK234RI10      eSK234        RI10               d3       Day_3           A
eSK239RI5       eSK239         RI5               d0       Day_0           A
eSK239RI6       eSK239         RI6               d3       Day_3           A
eSK239RI7       eSK239         RI7               d3       Day_3           A
eSK239RI8       eSK239         RI8               d3       Day_3           A
...                ...         ...              ...         ...         ...
eSK238RI11      eSK238        RI11               d9       Day_9           A
eSK238RI12      eSK238        RI12               d9       Day_9           A
eSK238RI13      eSK238        RI13               d9       Day_9           A
eSK238RI15      eSK238        RI15               d0       Day_0           A
eSK238RI16      eSK238        RI16               d0       Day_0           A
               sortday      digest
           <character> <character>
eSK234RI10           B           C
eSK239RI5            B           C
eSK239RI6            B           C
eSK239RI7            B           C
eSK239RI8            B           C
...                ...         ...
eSK238RI11           B           C
eSK238RI12           B           C
eSK238RI13           B           C
eSK238RI15           B           C
eSK238RI16           B           C
```

```r
table(prot$SampleType)
```

```
Day_0 Day_3 Day_9
  147   139   135
```

```
# Proportion of missing values
mean(is.na(assay(prot)))
```

```
[1] 0.791287
```

```
# Proportion of missing values in proteins that are missing in all cells of a TMT
 ↪  set
na_in_blocks <- sapply(split(colnames(prot), prot$Set), function(cells) {
    na <- is.na(assay(prot)[, cells, drop = FALSE])
    sum(rowSums(na) == length(cells)) * length(cells)
})
sum(na_in_blocks) / sum(is.na(assay(prot)))
```

```
[1] 0.9817978
```

About 80% of the values are missing, and nearly all missing values occur in proteins that are missing in all cells of a TMT set (i.e., an MS run). Thus, missingness is mostly determined by which peptides were selected for fragmentation and identified in each run, not by low abundances in individual cells (Vanderaa and Gatto 2023b). This has important implications for how we handle missing values. First, imputation methods that assume left-censored data (e.g., MinProb) are not suitable, as they would assign artificially low values to all cells of a set, creating spurious co-abundance among proteins that are missing in the same sets. This is aggravated by the fact that values are log-transformed *relative* abundances centered on each protein's mean, so that low values do not indicate low absolute abundances. Second, the data must be corrected for batch effects (i.e., differences between TMT sets). Hence, following current recommendations for SCP data analysis (Vanderaa and Gatto 2023b, 2025), we will avoid imputation whenever possible, using methods that handle missing values directly.

Let's add gene symbols to the `rowData`, since proteins are identified by UniProt accessions.

```
# Add gene symbols to `rowData`
acc <- sub("-.*", "", rownames(prot)) # remove isoform suffixes
rowData(prot)$symbol <- AnnotationDbi::mapIds(
    org.Hs.eg.db::org.Hs.eg.db, keys = acc, keytype = "UNIPROT",
    column = "SYMBOL", multiVals = "first"
)
```

#### Aggregating cells into metacells

To create metacells, we will first remove differences between TMT sets with `limma::removeBatchEffect()` (Ritchie et al. 2015), while preserving differences between time points. Since **limma** fits linear models to each protein using only observed values, missing values are kept as they are. Then, we will identify groups of similar cells with a principal component analysis (PCA) using the NIPALS algorithm (as implemented in the **nipals** package (Wright 2026)), which handles missing values without imputation (Vanderaa and Gatto 2025). To avoid restricting the analysis to the most consistently detected (and, hence, highly abundant) proteins, we will use all proteins detected in at least 30% of the cells.

```
# Remove differences between TMT sets, preserving differences between days
mat <- limma::removeBatchEffect(
    assay(prot), batch = prot$Set, design = model.matrix(~ prot$SampleType)
)

# PCA with NIPALS using proteins detected in at least 30% of the cells
na_prop <- rowMeans(is.na(mat))
sum(na_prop <= 0.7)
```

```
[1] 876
```

```r
pca <- nipals::nipals(t(mat[na_prop <= 0.7, ]), ncomp = 10, scale = FALSE)
pcs <- pca$scores

# Proportion of variance in the top 3 PCs explained by days and TMT sets
sapply(c(day = "SampleType", set = "Set"), function(covariate) {
    sapply(1:3, function(k) summary(lm(pcs[, k] ~ prot[[covariate]]))$r.squared)
})
```

```
            day         set
[1,] 0.77098150 0.043673608
[2,] 0.55986799 0.020346641
[3,] 0.03775828 0.002696338
```

The top principal components reflect differences between time points, not between TMT sets. Now, we will cluster cells from each time point with k-means, aiming at ~7 cells per metacell, and average the abundances of cells in each metacell (ignoring missing values).

```r
# Cluster cells within each time point into metacells
cells_by_day <- split(colnames(prot), prot$SampleType)
metacell <- unlist(lapply(names(cells_by_day), function(day) {
    cells <- cells_by_day[[day]]
    km <- kmeans(pcs[cells, ], centers = round(length(cells) / 7), nstart = 10)
    setNames(paste0(day, "_mc", km$cluster), cells)
}))[colnames(prot)]

# Number of cells and TMT sets per metacell
summary(as.vector(table(metacell)))
```

```
   Min. 1st Qu.  Median    Mean 3rd Qu.    Max. 
  1.000   4.000   6.000   7.017  10.000  16.000 
```

```r
summary(as.vector(tapply(prot$Set, metacell, function(s) length(unique(s)))))
```

```
   Min. 1st Qu.  Median    Mean 3rd Qu.    Max. 
  1.000   4.000   6.000   6.283   9.000  14.000 
```

```r
# Average abundances in each metacell
mc_mat <- sapply(split(seq_along(metacell), metacell), function(i) {
    rowMeans(mat[, i, drop = FALSE], na.rm = TRUE)
})
mc_mat[is.nan(mc_mat)] <- NA
dim(mc_mat)
```

```
[1] 4096   60
```

```r
mean(is.na(mc_mat))
```

```
[1] 0.5746053
```

Since metacells contain cells from different TMT sets, aggregation substantially reduced the proportion of missing values. Let's see how this affects correlations between proteins (using only proteins with few missing values, so that correlations can be estimated for cells).

```r
# Distribution of absolute correlations for proteins with few missing values
p <- na_prop <= 0.1
cor_cells <- cor(t(mat[p, ]), use = "pairwise.complete.obs")
cor_mc <- cor(t(mc_mat[p, ]), use = "pairwise.complete.obs")

data.frame(
    level = rep(c("Cells", "Metacells"), each = sum(upper.tri(cor_cells))),
    abs_cor = abs(c(cor_cells[upper.tri(cor_cells)], cor_mc[upper.tri(cor_mc)]))
) |>
    group_by(level) |>
    summarise(mean = mean(abs_cor), median = median(abs_cor))
```

```
# A tibble: 2 x 3
  level      mean median
  <chr>     <dbl>  <dbl>
1 Cells     0.164  0.131
2 Metacells 0.273  0.240
```

Absolute correlations increased substantially after aggregation, indicating that sparsity was masking co-abundance signals. For network inference, we will keep proteins detected in at least 80% of the metacells, as correlations estimated from few observations are unreliable. Since correlations in **BioNERO** are calculated from complete data, we will replace the remaining missing values (a small fraction) with protein means using `BioNERO::replace_na()`. Unlike left-censored imputation, mean imputation does not introduce artificial structure in the data, and it can only shrink correlations toward zero (i.e., it is conservative). Then, we will create a `GraphExperiment` object with metacells in columns, and store the mapping between cells and metacells in the `metadata` slot.

```r
# Keep proteins detected in >=80% of the metacells and replace NAs
mc_mat <- mc_mat[rowMeans(is.na(mc_mat)) <= 0.2, ]
mean(is.na(mc_mat))
```

```
[1] 0.04530142
```

```r
mc_mat <- BioNERO::replace_na(mc_mat, replaceby = "mean")

# Create metacell metadata
mc_md <- data.frame(
    row.names = colnames(mc_mat),
    day = sub("_mc.*", "", colnames(mc_mat)),
    n_cells = as.vector(table(metacell)[colnames(mc_mat)])
)

# Create `GraphExperiment` object
ge <- GraphExperiment(
    assays = list(abundance = mc_mat),
    rowData = rowData(prot)[rownames(mc_mat), ],
    colData = mc_md,
    metadata = list(metacells = data.frame(cell = colnames(prot), metacell))
)

ge
```

```
class: GraphExperiment
dim: 940 60
metadata(1): metacells
assays(1): abundance
```

```
rownames(940): A0A0U1RQI7 A1A4S6 ... Q9Y6N5 Q9Y6X9-2
rowData names(2): protein symbol
colnames(60): Day_0_mc1 Day_0_mc10 ... Day_9_mc8 Day_9_mc9
colData names(2): day n_cells
reducedDimNames(0):
mainExpName: NULL
altExpNames(0):
rowGraphs(0):
colGraphs(0):
```

#### Inferring a protein co-abundance network

Now, we will infer a signed hybrid co-abundance network with **BioNERO** (Almeida-Silva and Venancio 2021). As in use case 1, we will store the full network (i.e., a complete weighted graph) in `rowGraphs`, with module assignments and intramodular degrees as node attributes.

```
# Get beta power
sft <- SFT_fit(ge, net_type = "signed hybrid", cor_method = "pearson")
```

```
   Power SFT.R.sq slope truncated.R.sq mean.k. median.k. max.k.
1      3    0.732 -1.16          0.964 13.5000  1.12e+01  44.50
2      4    0.868 -1.23          0.983  6.9200  4.90e+00  29.70
3      5    0.896 -1.25          0.972  4.0100  2.31e+00  21.20
4      6    0.916 -1.34          0.978  2.5300  1.15e+00  16.80
5      7    0.918 -1.35          0.982  1.7000  5.99e-01  13.60
6      8    0.929 -1.36          0.977  1.2000  3.27e-01  11.20
7      9    0.886 -1.38          0.908  0.8800  1.89e-01   9.32
8     10    0.909 -1.37          0.936  0.6640  1.11e-01   7.85
9     11    0.915 -1.33          0.935  0.5130  6.57e-02   6.67
10    12    0.876 -1.35          0.888  0.4050  3.95e-02   5.70
11    13    0.956 -1.28          0.975  0.3250  2.43e-02   4.91
12    14    0.964 -1.27          0.979  0.2650  1.49e-02   4.25
13    15    0.978 -1.22          0.985  0.2190  9.26e-03   3.69
14    16    0.973 -1.20          0.973  0.1820  5.86e-03   3.23
15    17    0.974 -1.18          0.969  0.1540  3.69e-03   2.83
16    18    0.974 -1.16          0.969  0.1300  2.32e-03   2.49
17    19    0.957 -1.15          0.945  0.1120  1.48e-03   2.20
18    20    0.962 -1.12          0.951  0.0964  9.59e-04   1.95
```

```
sft$power
```

```
[1] 4
```

```
# Infer co-abundance network
net <- exp2gcn(
    ge, net_type = "signed hybrid", SFTpower = sft$power,
    cor_method = "pearson"
)
```

```
..connectivity..
..matrix multiplication (system BLAS)..
..normalization..
..done.
```

```r
table(net$genes_and_modules$Modules)
```

```
      black        blue       brown       green greenyellow        grey
         64          30         126          45         166         249
    magenta        pink         red      yellow
         89          75          56          40
```

```r
# Create igraph object and add node attributes
g <- graph_from_adjacency_matrix(
    net$adjacency_matrix, mode = "undirected", weighted = TRUE, diag = FALSE
)
modules <- setNames(net$genes_and_modules$Modules, net$genes_and_modules$Genes)
V(g)$module <- modules[V(g)$name]
V(g)$kWithin <- net$kIN[V(g)$name, "kWithin"]

rowGraph(ge, "wgcna") <- g
```

We will also add a kNN graph of metacells to the `colGraphs` slot, which can be used, for instance, to visualize or cluster metacells.

```r
# Build kNN graph of metacells from top PCs
mc_pcs <- prcomp(t(assay(ge, "abundance")), rank. = 10)$x
g_mc <- makeKNNGraph(mc_pcs, k = 5)
V(g_mc)$name <- colnames(ge)

colGraph(ge, "metacell_knn") <- g_mc

ge
```

```
class: GraphExperiment
dim: 940 60
metadata(1): metacells
assays(1): abundance
rownames(940): A0A0U1RQI7 A1A4S6 ... Q9Y6N5 Q9Y6X9-2
rowData names(4): protein symbol wgcna__module wgcna__kWithin
colnames(60): Day_0_mc1 Day_0_mc10 ... Day_9_mc8 Day_9_mc9
colData names(2): day n_cells
reducedDimNames(0):
mainExpName: NULL
altExpNames(0):
rowGraphs(1): wgcna
colGraphs(1): metacell_knn
```

#### Exploring the co-abundance network

Module assignments are now available in `rowData`, and time points are in `colData`. Thus, we can easily identify modules associated with EMT stages by correlating module eigengenes with time points.

```r
# Correlate module eigengenes with time points
me_trait <- module_trait_cor(ge, MEs = net$MEs, metadata_cols = "day")
plot_module_trait_cor(me_trait)
```

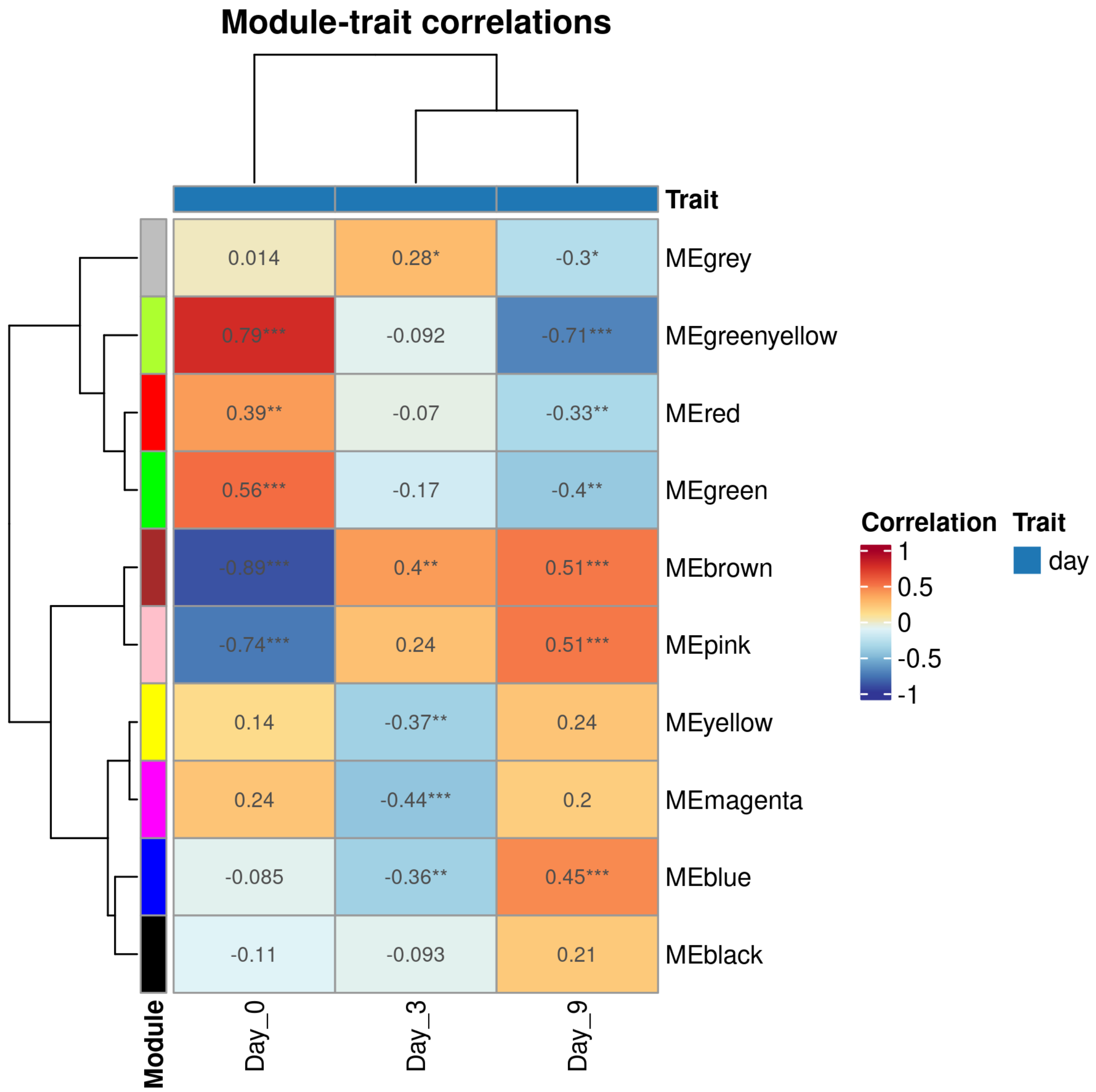

Let's look at the top hub proteins of the module most negatively associated with day 0 (i.e., proteins with increased abundances along EMT).

```
# Get module most negatively correlated with day 0
emt_module <- me_trait |>
    filter(trait == "Day_0") |>
    slice_min(cor, n = 1) |>
    pull(ME) |>
    str_remove("^ME")
emt_module
```

```
[1] "brown"
```

```
# Top 20 proteins with the highest intramodular degree in this module
top_proteins <- rowData(ge) |>
    as.data.frame() |>
    rownames_to_column("accession") |>
    filter(wgcna__module == emt_module) |>
    slice_max(wgcna__kWithin, n = 20)

top_proteins$symbol
```

```
 [1] "MYL6"    "MYH9"    "MSN"     "ACTB"    "TP53I3"  "CNN2"    "CECR2"
 [8] "FLNB"    "TPM3"    "TAGLN2"  "TPM4"    "KLHDC7B" "TGM2"    "ARPC4"
[15] "RRBP1"   "MYL12A"  "CFL1"    "TAGLN"   "ACTR2"   "FLNA"
```

Top hubs include several proteins involved in actomyosin contractility and cytoskeletal remodeling (e.g., myosins, tropomyosins, filamins, and transgelins), a hallmark of the transition to a mesenchymal phenotype.

Finally, let's visualize the top hubs of this module. Since `rowGraph()` adds `rowData` variables as node attributes, gene symbols are readily available for labeling.

```r
# Subset object to keep top hubs and extract graph
g_hubs <- rowGraph(ge[top_proteins$accession, ], "wgcna")

# For visualization, keep only the strongest edges
g_hubs <- delete_edges(
    g_hubs, which(E(g_hubs)$weight < quantile(E(g_hubs)$weight, 0.5))
)

ggraph(g_hubs, layout = "fr") +
    geom_edge_link(aes(width = weight), alpha = 0.3, color = "grey50") +
    geom_node_point(aes(size = kWithin), color = emt_module) +
    geom_node_text(aes(label = symbol), repel = TRUE, size = 3.5) +
    scale_edge_width(range = c(0.2, 2)) +
    theme_graph(base_family = "sans") +
    labs(size = "Intramodular\ndegree", edge_width = "Adjacency")
```

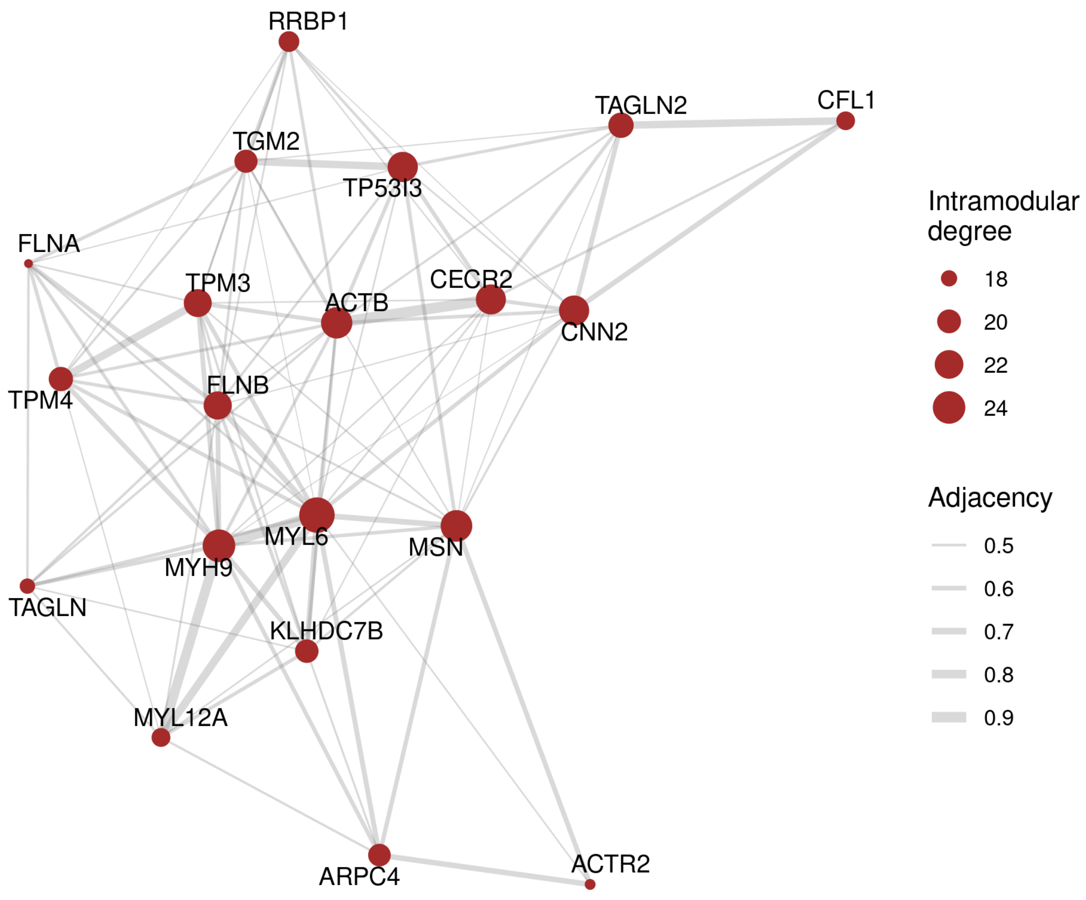

### Session info

This document was created under the following conditions:

```
- Session info ---------------------------------------------------------------
 setting  value
 version  R version 4.5.2 (2025-10-31)
 os       Ubuntu 22.04.5 LTS
 system   x86_64, linux-gnu
 ui       X11
 language (EN)
 collate  en_US.UTF-8
 ctype    en_US.UTF-8
 tz       Europe/Brussels
 date     2026-10-24
 pandoc   3.6.3 @ /opt/quarto/bin/tools/ (via rmarkdown)
 quarto   1.8.25 @ /usr/local/bin/quarto

- Packages -------------------------------------------------------------------
 package              * version date (UTC) lib source
 AnnotationHub        * 4.0.0   2025-10-29 [1] Bioconductor 3.22 (R 4.5.2)
 Biobase              * 2.70.0  2025-10-29 [1] Bioconductor 3.22 (R 4.5.2)
 BiocFileCache        * 3.0.0   2025-10-29 [1] Bioconductor 3.22 (R 4.5.2)
 BiocGenerics         * 0.56.0  2025-10-29 [1] Bioconductor 3.22 (R 4.5.2)
 BioNERO              * 1.18.0  2025-10-29 [1] Bioconductor 3.22 (R 4.5.2)
 bluster              * 1.20.0  2025-10-29 [1] Bioconductor 3.22 (R 4.5.2)
 dbplyr               * 2.5.1   2025-09-10 [1] CRAN (R 4.5.2)
 dplyr                * 1.1.4   2023-11-17 [1] CRAN (R 4.5.2)
 ExperimentHub        * 3.0.0   2025-10-29 [1] Bioconductor 3.22 (R 4.5.2)
 forcats              * 1.0.1   2025-09-25 [1] CRAN (R 4.5.2)
 generics             * 0.1.4   2025-05-09 [1] CRAN (R 4.5.2)
 GenomicRanges        * 1.62.1  2025-12-08 [1] Bioconductor 3.22 (R 4.5.2)
 ggplot2              * 4.0.1   2025-11-14 [1] CRAN (R 4.5.2)
 ggraph               * 2.2.2   2025-08-24 [1] CRAN (R 4.5.2)
 GraphExperiment      * 1.1.2   2026-09-08 [1] Bioconductor
 here                 * 1.0.2   2025-09-15 [1] CRAN (R 4.5.2)
 igraph               * 2.2.1   2025-10-27 [1] CRAN (R 4.5.2)
 IRanges              * 2.44.0  2025-10-29 [1] Bioconductor 3.22 (R 4.5.2)
 lubridate            * 1.9.4   2024-12-08 [1] CRAN (R 4.5.2)
 MatrixGenerics       * 1.22.0  2025-10-29 [1] Bioconductor 3.22 (R 4.5.2)
 matrixStats          * 1.5.0   2025-01-07 [1] CRAN (R 4.5.2)
 MultiAssayExperiment * 1.36.2  2026-03-29 [1] Bioconductor 3.22 (R 4.5.2)
 purrr                * 1.2.1   2026-01-09 [1] CRAN (R 4.5.2)
 QFeatures            * 1.20.0  2025-10-29 [1] Bioconductor 3.22 (R 4.5.2)
 readr                * 2.1.6   2025-11-14 [1] CRAN (R 4.5.2)
 S4Vectors            * 0.48.0  2025-10-29 [1] Bioconductor 3.22 (R 4.5.2)
 scpdata              * 1.18.0  2025-11-04 [1] Bioconductor 3.22 (R 4.5.2)
 Seqinfo              * 1.0.0   2025-10-29 [1] Bioconductor 3.22 (R 4.5.2)
 SingleCellExperiment * 1.32.0  2025-10-29 [1] Bioconductor 3.22 (R 4.5.2)
 stringr              * 1.6.0   2025-11-04 [1] CRAN (R 4.5.2)
 SummarizedExperiment * 1.40.0  2025-10-29 [1] Bioconductor 3.22 (R 4.5.2)
 tibble               * 3.3.1   2026-01-11 [1] CRAN (R 4.5.2)
 tidyr                * 1.3.2   2025-12-19 [1] CRAN (R 4.5.2)
 tidyverse            * 2.0.0   2023-02-22 [1] CRAN (R 4.5.2)

 [1] /home/faalm/R/x86_64-pc-linux-gnu-library/4.5
 [2] /usr/local/lib/R/site-library
 [3] /usr/lib/R/site-library
 [4] /usr/lib/R/library
```

```
 * -- Packages attached to the search path.

------------------------------------------------------------------------------
```

### Use case 3: microbial co-occurrence networks

Here, we will use **GraphExperiment** to store microbial association networks inferred from 16S rRNA amplicon sequencing data, with taxon-taxon networks in `rowGraphs` and a sample-sample network in `colGraphs`. We will use the `peerj13075` data set from the **mia** package, a core package of the miaverse ecosystem (Borman et al. 2025), which contains skin microbiota profiles of 58 genetically unrelated Indian individuals from three different cities and with different diets (vegetarian and mixed) (Potbhare et al. 2022).

Microbiome abundance data are *compositional*, i.e., sequencing only provides information on the relative abundances of taxa. Thus, standard correlation measures (e.g., Pearson's or Spearman's correlations) can lead to spurious associations (Friedman and Alm 2012; Gloor et al. 2017). To account for compositionality, we will use two widely used methods implemented in the **SpiecEasi** package: SPIEC-EASI (Kurtz et al. 2015), which infers a sparse graphical model (i.e., conditional dependencies between taxa) from centered log-ratio (CLR)-transformed data, and SparCC (Friedman and Alm 2012), which estimates correlations between the underlying (log-transformed) absolute abundances.

```
library(here)
library(GraphExperiment)
library(mia)
library(scater)
library(SpiecEasi)
library(bluster)
library(tidyverse)
library(ggraph)
library(patchwork)

set.seed(123) # for reproducibility
```

#### Data description and processing

The data are stored in a `TreeSummarizedExperiment` object, the core data structure of the miaverse ecosystem (Huang et al. 2020). Taxa were already agglomerated at the genus level.

```
# Load data
data("peerj13075", package = "mia")
tse <- peerj13075
tse
```

```
class: TreeSummarizedExperiment
dim: 674 58
metadata(0):
assays(1): counts
rownames(674): OTU1 OTU2 ... OTU2567 OTU2569
rowData names(6): kingdom phylum ... family genus
colnames(58): ID1 ID2 ... ID57 ID58
colData names(5): Sample Geographical_location Gender Age Diet
reducedDimNames(0):
mainExpName: NULL
altExpNames(0):
rowLinks: NULL
rowTree: NULL
```

```
colLinks: NULL
colTree: NULL
```

```r
# Sample metadata
table(tse$Geographical_location, tse$Diet)
```

```
             Mixed Veg
  Ahmednagar     4   7
  Nashik        10   1
  Pune          14  22
```

Since rare taxa provide little information for network inference, we will only keep genera with at least one read in at least 20% of the samples. Then, we will add a CLR-transformed assay, which we will use later.

```r
# Keep only prevalent genera
tse <- subsetByPrevalent(tse, detection = 1, prevalence = 0.2)
dim(tse)
```

```
[1] 72 58
```

```r
# Add CLR-transformed assay
tse <- transformAssay(tse, method = "clr", pseudocount = 1)
```

Like `GraphExperiment`, `TreeSummarizedExperiment` extends the `SingleCellExperiment` class. Hence, we can easily coerce a `TreeSummarizedExperiment` object to a `GraphExperiment` object. Tree-specific slots (`rowTree`, `colTree`, etc.) are not kept, but they could be stored in the `metadata` slot if needed (see use cases 6 and 7 for examples on how to handle phylogenies).

```r
# Coerce `TreeSummarizedExperiment` to `GraphExperiment`
ge <- as(tse, "GraphExperiment")
ge
```

```
class: GraphExperiment
dim: 72 58
metadata(0):
assays(2): counts clr
rownames(72): OTU2 OTU53 ... OTU2544 OTU2550
rowData names(6): kingdom phylum ... family genus
colnames(58): ID1 ID2 ... ID57 ID58
colData names(5): Sample Geographical_location Gender Age Diet
reducedDimNames(0):
mainExpName: NULL
altExpNames(0):
rowGraphs(0):
colGraphs(0):
```

#### Inferring taxon-taxon association networks

Now, we will infer an association network with SPIEC-EASI using the Meinshausen-Bühlmann neighborhood selection method, with the sparsity parameter selected with the StARS approach (Liu, Roeder, and Wasserman 2010). Note that `spiec.easi()` expects samples in rows and taxa in columns, and it performs CLR transformation internally.

```r
# Infer network with SPIEC-EASI
counts <- t(assay(ge, "counts"))
se_mb <- spiec.easi(
    counts, method = "mb", lambda.min.ratio = 1e-3, nlambda = 30,
    pulsar.params = list(rep.num = 50, seed = 123)
)

# Get edge weights (symmetrized regression coefficients)
mb_weights <- as.matrix(symBeta(getOptBeta(se_mb), mode = "maxabs"))
dimnames(mb_weights) <- list(colnames(counts), colnames(counts))

# Create igraph object
g_spieceasi <- graph_from_adjacency_matrix(
    mb_weights, mode = "undirected", weighted = TRUE, diag = FALSE
)
E(g_spieceasi)$sign <- ifelse(E(g_spieceasi)$weight > 0, "positive", "negative")

g_spieceasi
```

```
IGRAPH 958d388 UNW- 72 61 --
+ attr: name (v/c), weight (e/n), sign (e/c)
+ edges from 958d388 (vertex names):
 [1] OTU53  --OTU978  OTU119 --OTU520  OTU119 --OTU929  OTU119 --OTU2111
 [5] OTU119 --OTU2377 OTU194 --OTU773  OTU194 --OTU1768 OTU198 --OTU2243
 [9] OTU367 --OTU1075 OTU520 --OTU2076 OTU520 --OTU2111 OTU562 --OTU1075
[13] OTU562 --OTU1585 OTU804 --OTU929  OTU804 --OTU2054 OTU804 --OTU2076
[17] OTU807 --OTU1212 OTU860 --OTU1075 OTU860 --OTU1622 OTU860 --OTU1810
[21] OTU860 --OTU2059 OTU860 --OTU2529 OTU929 --OTU2076 OTU978 --OTU987
[25] OTU978 --OTU1709 OTU987 --OTU1709 OTU987 --OTU1789 OTU987 --OTU2076
[29] OTU987 --OTU2111 OTU987 --OTU2371 OTU1034--OTU1768 OTU1075--OTU1336
+ ... omitted several edges
```

Next, we will estimate SparCC correlations, and use 100 bootstrap replicates to obtain P-values. We will keep only edges with P $< 0.05$ and absolute correlations $\geq 0.3$.

```r
# Estimate SparCC correlations with bootstrap P-values
sparcc_boot <- sparccboot(counts, R = 100)
sparcc_p <- pval.sparccboot(sparcc_boot)

# Create matrices of correlations and P-values
n <- ncol(counts)
sparcc_cor <- sparcc_pval <- matrix(
    0, n, n, dimnames = list(colnames(counts), colnames(counts))
)
sparcc_cor[upper.tri(sparcc_cor)] <- sparcc_p$cors
sparcc_pval[upper.tri(sparcc_pval)] <- sparcc_p$pvals

# Keep only significant correlations
sparcc_cor[sparcc_pval >= 0.05 | abs(sparcc_cor) < 0.3] <- 0

# Create igraph object
g_sparcc <- graph_from_adjacency_matrix(
    sparcc_cor, mode = "upper", weighted = TRUE, diag = FALSE
)
E(g_sparcc)$sign <- ifelse(E(g_sparcc)$weight > 0, "positive", "negative")

g_sparcc
```

```
IGRAPH 363b598 UNW- 72 249 --
+ attr: name (v/c), weight (e/n), sign (e/c)
+ edges from 363b598 (vertex names):
 [1] OTU53 --OTU119  OTU53 --OTU520  OTU53 --OTU804  OTU53 --OTU929
 [5] OTU53 --OTU978  OTU53 --OTU987  OTU53 --OTU1585 OTU53 --OTU1680
 [9] OTU53 --OTU1709 OTU53 --OTU1810 OTU53 --OTU2076 OTU53 --OTU2377
[13] OTU53 --OTU2529 OTU119--OTU520  OTU119--OTU804  OTU119--OTU929
[17] OTU119--OTU978  OTU119--OTU987  OTU119--OTU1260 OTU119--OTU1585
[21] OTU119--OTU1680 OTU119--OTU1709 OTU119--OTU2076 OTU119--OTU2111
[25] OTU119--OTU2117 OTU119--OTU2127 OTU119--OTU2243 OTU119--OTU2371
[29] OTU119--OTU2377 OTU194--OTU540  OTU194--OTU773  OTU194--OTU1034
+ ... omitted several edges
```

Both networks were inferred from the same features, so they can be stored together in the `rowGraphs` slot. We will also add node degrees as node attributes.

```r
# Add node degree as node attributes
V(g_spieceasi)$degree <- degree(g_spieceasi)
V(g_sparcc)$degree <- degree(g_sparcc)

# Add graphs to `rowGraphs`
rowGraphs(ge) <- list(spieceasi = g_spieceasi, sparcc = g_sparcc)

ge
```

```
class: GraphExperiment
dim: 72 58
metadata(0):
assays(2): counts clr
rownames(72): OTU2 OTU53 ... OTU2544 OTU2550
rowData names(8): kingdom phylum ... spieceasi__degree sparcc__degree
colnames(58): ID1 ID2 ... ID57 ID58
colData names(5): Sample Geographical_location Gender Age Diet
reducedDimNames(0):
mainExpName: NULL
altExpNames(0):
rowGraphs(2): spieceasi sparcc
colGraphs(0):
```

#### Inferring a sample-sample network

Microbiome data are often explored at the sample level, e.g., to identify groups of samples with similar community compositions. Here, we will create a k-nearest neighbor (kNN) graph of samples based on Aitchison distances (i.e., Euclidean distances of CLR-transformed data) with **bluster** (A. Lun 2025). For that, we will first perform a principal component analysis on the CLR-transformed data with **scater** (McCarthy et al. 2017), and then build the kNN graph from the top 10 principal components. Then, we will identify communities in this graph with the Louvain algorithm, and store community assignments as node attributes.

```r
# Perform PCA on CLR-transformed data
ge <- runPCA(ge, assay.type = "clr", ncomponents = 10)

# Build kNN graph of samples
g_samples <- makeKNNGraph(reducedDim(ge, "PCA"), k = 5)
V(g_samples)$name <- colnames(ge)

# Identify communities of samples
V(g_samples)$community <- as.character(membership(cluster_louvain(g_samples)))
```

```r
# Add graph to `colGraphs`
colGraph(ge, "aitchison_knn") <- g_samples

ge
```

```
class: GraphExperiment
dim: 72 58
metadata(0):
assays(2): counts clr
rownames(72): OTU2 OTU53 ... OTU2544 OTU2550
rowData names(8): kingdom phylum ... spieceasi__degree sparcc__degree
colnames(58): ID1 ID2 ... ID57 ID58
colData names(6): Sample Geographical_location ... Diet
  aitchison_knn__community
reducedDimNames(1): PCA
mainExpName: NULL
altExpNames(0):
rowGraphs(2): spieceasi sparcc
colGraphs(1): aitchison_knn
```

Note that the `GraphExperiment` object now contains assays, taxon and sample metadata, principal components, and graphs of both taxa and samples. As node attributes are added to `rowData` and `colData`, we can explore how communities of samples are associated with sample metadata.

```r
# Community assignments vs sample metadata
table(ge$aitchison_knn__community, ge$Geographical_location)
```

```
    Ahmednagar Nashik Pune
  1          2      1    8
  2          5      0    1
  3          2      1    9
  4          0      0    5
  5          0      1    8
  6          0      7    0
  7          2      1    5
```

```r
table(ge$aitchison_knn__community, ge$Diet)
```

```
    Mixed Veg
  1     5   6
  2     2   4
  3     3   9
  4     2   3
  5     4   5
  6     7   0
  7     5   3
```

#### Exploring and comparing taxon-taxon networks

Since both networks are stored in the same object, we can easily compare them. For instance, let's check how many edges are shared between the networks, and whether shared edges have the same sign.

```r
# Get edges shared between networks
shared <- intersection(
    rowGraph(ge, "spieceasi"), rowGraph(ge, "sparcc"), keep.all.vertices = FALSE
)

c(
    SPIEC_EASI = ecount(rowGraph(ge, "spieceasi")),
    SparCC = ecount(rowGraph(ge, "sparcc")),
    Shared = ecount(shared)
)
```

```
SPIEC_EASI     SparCC     Shared
        61        249         51
```

```r
# Same sign in both networks?
table(E(shared)$sign_1 == E(shared)$sign_2)
```

```
TRUE
  51
```

Node degrees in each graph are available in `rowData`, along with taxonomic information. Thus, we can identify the most connected taxa (hubs) in each network, and check whether they agree.

```r
rowData(ge) |>
    as.data.frame() |>
    select(phylum, genus, spieceasi__degree, sparcc__degree) |>
    arrange(-spieceasi__degree) |>
    head(10)
```

```
                phylum                genus spieceasi__degree sparcc__degree
OTU2529     Firmicutes        Virgibacillus                 9             22
OTU2111 Proteobacteria             Serratia                 8             19
OTU987  Proteobacteria Escherichia/Shigella                 6             10
OTU2076 Proteobacteria           Salmonella                 6             20
OTU860      Firmicutes     Desulfosporomusa                 5             13
OTU1075     Firmicutes          Geobacillus                 5             20
OTU119  Proteobacteria            Aeromonas                 4             17
OTU1622     Firmicutes    Ornithinibacillus                 4             10
OTU1810     Firmicutes        Planomicrobium                 4             13
OTU520      Firmicutes        Brevibacillus                 3             23
```

```r
# Are degrees correlated?
cor.test(
    rowData(ge)$spieceasi__degree, rowData(ge)$sparcc__degree,
    method = "spearman", exact = FALSE
)
```

```
    Spearman's rank correlation rho

data:  rowData(ge)$spieceasi__degree and rowData(ge)$sparcc__degree
S = 13359, p-value = 3.226e-16
alternative hypothesis: true rho is not equal to 0
sample estimates:
      rho
0.7852122
```

Finally, let's visualize both networks with nodes colored by phylum. Since `rowGraph()` adds `rowData` variables as node attributes, taxonomic information is readily available for plotting.

```
# Define colors for phyla (only those of connected nodes)
connected <- rowData(ge)$spieceasi__degree > 0 | rowData(ge)$sparcc__degree > 0
phyla <- sort(unique(rowData(ge)$phylum[connected]))
phylum_colors <- setNames(palette.colors(length(phyla), "Okabe-Ito"), phyla)
max_degree <- max(rowData(ge)$spieceasi__degree, rowData(ge)$sparcc__degree)

# Plotting function
plot_network <- function(g, title) {
    g <- delete_vertices(g, which(degree(g) == 0))
    ggraph(g, layout = "fr", weights = NA) +
        geom_edge_link(aes(color = sign), alpha = 0.6) +
        geom_node_point(aes(fill = phylum, size = degree), shape = 21) +
        scale_edge_color_manual(
            values = c(positive = "#009E73", negative = "#D55E00")
        ) +
        scale_fill_manual(values = phylum_colors, limits = phyla) +
        scale_size_continuous(range = c(1.5, 6), limits = c(1, max_degree)) +
        theme_graph(base_family = "sans") +
        labs(title = title, fill = "Phylum", size = "Degree", edge_color =
         ↪  "Association")
}

# Plot networks
wrap_plots(
    plot_network(rowGraph(ge, "spieceasi"), "SPIEC-EASI (MB)"),
    plot_network(rowGraph(ge, "sparcc"), "SparCC"),
    nrow = 1
) +
    plot_layout(guides = "collect") +
    plot_annotation(tag_levels = "A") &
    theme(plot.title = element_text(size = 13))
```

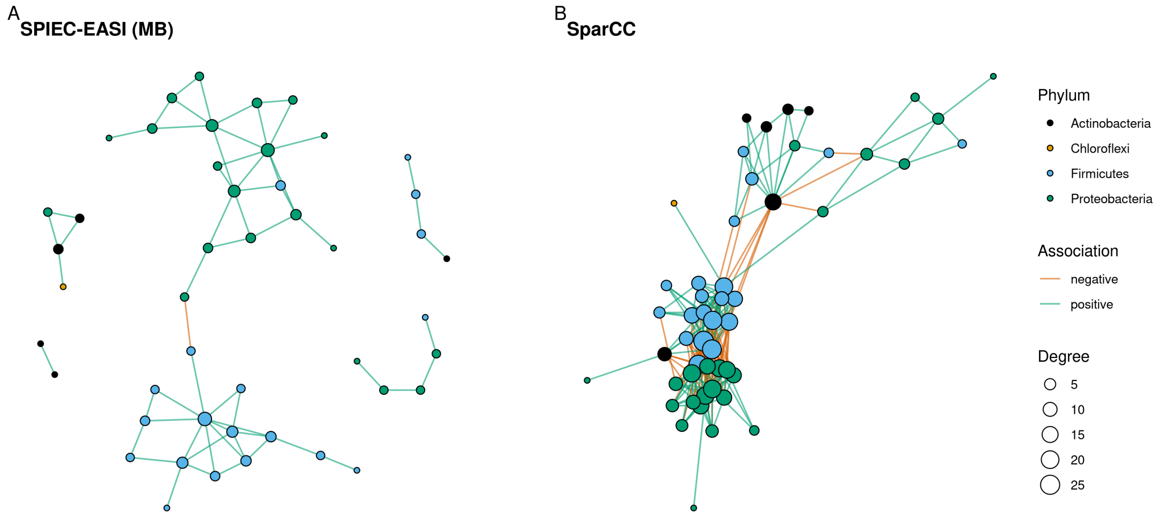

Other compositionally-aware methods (e.g., SPRING, CCLasso, or proportionality-based measures), as well as tools for network comparison, are available in the **NetCoMi** package (Peschel et al. 2020). Regardless of the method used, the resulting networks can be stored in the same `GraphExperiment` object, making it easy to compare and integrate them.

#### Session info

This document was created under the following conditions:

```
- Session info ---------------------------------------------------------------
 setting  value
 version  R version 4.5.2 (2025-10-31)
 os       Ubuntu 22.04.5 LTS
 system   x86_64, linux-gnu
 ui       X11
 language (EN)
 collate  en_US.UTF-8
 ctype    en_US.UTF-8
 tz       Europe/Brussels
 date     2026-10-24
 pandoc   3.6.3 @ /opt/quarto/bin/tools/ (via rmarkdown)
 quarto   1.8.25 @ /usr/local/bin/quarto

- Packages -------------------------------------------------------------------
 package                  * version date (UTC) lib source
 Biobase                  * 2.70.0  2025-10-29 [1] Bioconductor 3.22 (R 4.5.2)
 BiocGenerics             * 0.56.0  2025-10-29 [1] Bioconductor 3.22 (R 4.5.2)
 Biostrings               * 2.78.0  2025-10-29 [1] Bioconductor 3.22 (R 4.5.2)
 bluster                  * 1.20.0  2025-10-29 [1] Bioconductor 3.22 (R 4.5.2)
 dplyr                    * 1.1.4   2023-11-17 [1] CRAN (R 4.5.2)
 forcats                  * 1.0.1   2025-09-25 [1] CRAN (R 4.5.2)
 generics                 * 0.1.4   2025-05-09 [1] CRAN (R 4.5.2)
 GenomicRanges            * 1.62.1  2025-12-08 [1] Bioconductor 3.22 (R 4.5.2)
 ggplot2                  * 4.0.1   2025-11-14 [1] CRAN (R 4.5.2)
 ggraph                   * 2.2.2   2025-08-24 [1] CRAN (R 4.5.2)
 GraphExperiment          * 1.1.2   2026-09-08 [1] Bioconductor
 here                     * 1.0.2   2025-09-15 [1] CRAN (R 4.5.2)
 igraph                   * 2.2.1   2025-10-27 [1] CRAN (R 4.5.2)
 IRanges                  * 2.44.0  2025-10-29 [1] Bioconductor 3.22 (R 4.5.2)
 lubridate                * 1.9.4   2024-12-08 [1] CRAN (R 4.5.2)
 MatrixGenerics           * 1.22.0  2025-10-29 [1] Bioconductor 3.22 (R 4.5.2)
 matrixStats              * 1.5.0   2025-01-07 [1] CRAN (R 4.5.2)
 mia                      * 1.18.0  2025-10-29 [1] Bioconductor 3.22 (R 4.5.2)
 MultiAssayExperiment     * 1.36.2  2026-03-29 [1] Bioconductor 3.22 (R 4.5.2)
 patchwork                * 1.3.2   2025-08-25 [1] CRAN (R 4.5.2)
 purrr                    * 1.2.1   2026-01-09 [1] CRAN (R 4.5.2)
 readr                    * 2.1.6   2025-11-14 [1] CRAN (R 4.5.2)
 S4Vectors                * 0.48.0  2025-10-29 [1] Bioconductor 3.22 (R 4.5.2)
 scater                   * 1.38.1  2026-03-20 [1] Bioconductor 3.22 (R 4.5.2)
 scuttle                  * 1.20.0  2025-10-30 [1] Bioconductor 3.22 (R 4.5.2)
 Seqinfo                  * 1.0.0   2025-10-29 [1] Bioconductor 3.22 (R 4.5.2)
 SingleCellExperiment     * 1.32.0  2025-10-29 [1] Bioconductor 3.22 (R 4.5.2)
 SpiecEasi                * 1.99.0  2026-10-21 [1] Github (zdk123/SpiecEasi@03b96da)
 stringr                  * 1.6.0   2025-11-04 [1] CRAN (R 4.5.2)
 SummarizedExperiment     * 1.40.0  2025-10-29 [1] Bioconductor 3.22 (R 4.5.2)
 tibble                   * 3.3.1   2026-01-11 [1] CRAN (R 4.5.2)
 tidyr                    * 1.3.2   2025-12-19 [1] CRAN (R 4.5.2)
 tidyverse                * 2.0.0   2023-02-22 [1] CRAN (R 4.5.2)
 TreeSummarizedExperiment * 2.18.0  2025-10-30 [1] Bioconductor 3.22 (R 4.5.2)
 XVector                  * 0.50.0  2025-10-29 [1] Bioconductor 3.22 (R 4.5.2)

 [1] /home/faalm/R/x86_64-pc-linux-gnu-library/4.5
 [2] /usr/local/lib/R/site-library
 [3] /usr/lib/R/site-library
```

```
 [4] /usr/lib/R/library
 * -- Packages attached to the search path.

------------------------------------------------------------------------------
```

### Use case 4: cell-cell graphs for clustering single-cell data

Graph-based clustering is the standard approach to identify cell populations in single-cell RNA-seq (scRNA-seq) data. In this approach, a graph of cells is built from a low-dimensional representation of the data (e.g., top principal components), with each cell connected to its nearest neighbors, and then communities are detected in the graph (Amezquita et al. 2019). Although cell-cell graphs are central to scRNA-seq data analysis, they are typically discarded after clustering, and there is no dedicated container for them in the `SingleCellExperiment` class. Here, we will use **GraphExperiment** to store cell-cell graphs in the `colGraphs` slot along with the expression data and dimensionality reduction results. We will use the mouse brain data set from Zeisel et al. (2015), which includes ~3,000 cells from the mouse cortex and hippocampus, available in the **scRNAseq** package (Risso and Cole 2025).

```
library(here)
library(GraphExperiment)
library(scRNAseq)
library(scrapper)
library(bluster)
library(tidyverse)
library(ggraph)
library(patchwork)

set.seed(123) # for reproducibility
```

#### Data description and processing

We will first load the data and perform standard processing steps, including quality control, normalization, selection of highly variable genes (HVGs), and principal component analysis (PCA), using **scrapper** (A. T. L. Lun and Kancherla 2023), which provides fast implementations of common single-cell analysis steps.

```
# Load data
sce <- ZeiselBrainData()
sce
```

```
class: SingleCellExperiment
dim: 20006 3005
metadata(0):
assays(1): counts
rownames(20006): Tspan12 Tshz1 ... mt-Rnr1 mt-Nd4l
rowData names(1): featureType
colnames(3005): 1772071015_C02 1772071017_G12 ... 1772066098_A12
  1772058148_F03
colData names(9): tissue group # ... level1class level2class
reducedDimNames(0):
mainExpName: gene
altExpNames(2): repeat ERCC
```

```r
# Cell types (as annotated by the authors)
table(sce$level1class)
```

```
astrocytes_ependymal    endothelial-mural         interneurons 
                 224                  235                  290 
           microglia     oligodendrocytes        pyramidal CA1 
                  98                  820                  939 
        pyramidal SS 
                 399 
```

```r
# Quality control: remove low-quality cells based on library sizes,
# number of detected genes, and proportion of mitochondrial reads
is_mito <- rowData(sce)$featureType == "mito"
sce <- quickRnaQc.se(sce, subsets = list(mito = is_mito), filter.cells = TRUE)

# Normalization (library size factors) and log-transformation
sce <- normalizeRnaCounts.se(sce, size.factors = sce$sum)

# Feature selection and PCA
sce <- chooseRnaHvgs.se(sce, top = 2000)
sce <- runPca.se(sce, features = rowData(sce)$hvg, number = 25)

# Run t-SNE for visualization
sce <- runTsne.se(sce)
sce
```

```
class: SingleCellExperiment 
dim: 20006 2874 
metadata(2): qc PCA
assays(2): counts logcounts
rownames(20006): Tspan12 Tshz1 ... mt-Rnr1 mt-Nd4l
rowData names(6): featureType means ... residuals hvg
colnames(2874): 1772071015_C02 1772071017_G12 ... 1772063068_D01
  1772066098_A12
colData names(14): tissue group # ... keep sizeFactor
reducedDimNames(2): PCA TSNE
mainExpName: gene
altExpNames(2): repeat ERCC
```

#### Building cell-cell graphs

Now, we will build a shared nearest neighbor (SNN) graph from the top principal components with `scrapper::buildSnnGraph()`, in which two cells are connected if they share any of their k nearest neighbors, with edge weights defined by the rank of the shared neighbors (Xu and Su 2015). This function returns a list with the number of vertices, the edges, and their weights, which we will convert to an `igraph` object with cell IDs as node names. For comparison, we will also build a kNN graph with `bluster::makeKNNGraph()`.

Note that functions that perform graph-based clustering (e.g., `scrapper::clusterGraph.se()`) typically discard the graph or store it in the `metadata` slot, where it is not subset along with the cells. Using **GraphExperiment**, graphs are stored in a dedicated slot and kept in sync with the cells.

```r
# Build SNN graph and convert it to an `igraph` object
pcs <- reducedDim(sce, "PCA")
snn_list <- buildSnnGraph(t(pcs), num.neighbors = 10, weight.scheme = "ranked")
snn <- make_undirected_graph(snn_list$edges, n = snn_list$vertices)
```

```r
E(snn)$weight <- snn_list$weights
V(snn)$name <- colnames(sce)

# Build kNN graph
knn <- makeKNNGraph(pcs, k = 10)
V(knn)$name <- colnames(sce)

snn
```

```
IGRAPH bb065f5 UNW- 2874 82468 --
+ attr: name (v/c), weight (e/n)
+ edges from bb065f5 (vertex names):
 [1] 1772071015_C02--1772071017_G12 1772071015_C02--1772071017_A05
 [3] 1772071017_G12--1772071017_A05 1772071015_C02--1772071014_B06
 [5] 1772071017_G12--1772071014_B06 1772071017_A05--1772071014_B06
 [7] 1772071015_C02--1772067065_H06 1772071017_G12--1772067065_H06
 [9] 1772071014_B06--1772067065_H06 1772071017_A05--1772067065_H06
[11] 1772071015_C02--1772071017_E02 1772071017_G12--1772071017_E02
[13] 1772071017_A05--1772071017_E02 1772071014_B06--1772071017_E02
[15] 1772067065_H06--1772071017_E02 1772071015_C02--1772067065_B07
+ ... omitted several edges
```

Next, we will create a `GraphExperiment` object by coercing the `SingleCellExperiment` object and adding both graphs to the `colGraphs` slot.

```r
# Create `GraphExperiment` object
ge <- as(sce, "GraphExperiment")
colGraphs(ge) <- list(snn = snn, knn = knn)

ge
```

```
class: GraphExperiment
dim: 20006 2874
metadata(2): qc PCA
assays(2): counts logcounts
rownames(20006): Tspan12 Tshz1 ... mt-Rnr1 mt-Nd4l
rowData names(6): featureType means ... residuals hvg
colnames(2874): 1772071015_C02 1772071017_G12 ... 1772063068_D01
  1772066098_A12
colData names(14): tissue group # ... keep sizeFactor
reducedDimNames(2): PCA TSNE
mainExpName: gene
altExpNames(2): repeat ERCC
rowGraphs(0):
colGraphs(2): snn knn
```

#### Graph-based clustering

We will now detect communities in each graph with the Louvain algorithm. Community assignments will be added as node attributes, so that they are automatically available in `colData`. We will also add node strength (sum of edge weights) as a node attribute, since it is a measure of how well connected a cell is to other cells.

```r
# Detect communities in each graph and add as node attributes
colGraphs(ge) <- lapply(colGraphs(ge), function(graph) {
    V(graph)$cluster <- as.character(membership(cluster_louvain(graph)))
```

```r
    V(graph)$strength <- strength(graph)
    graph
})

# Community assignments and node strengths are now in `colData`
colData(ge)[, grepl("__", names(colData(ge)))]
```

```
DataFrame with 2874 rows and 4 columns
               snn__cluster snn__strength knn__cluster knn__strength
                <character>     <numeric>  <character>     <numeric>
1772071015_C02            1         278.0            1            17
1772071017_G12            1         227.0            1            10
1772071017_A05            1         309.5            1            16
1772071014_B06            1         222.5            1            10
1772067065_H06            1         219.5            1            14
...                     ...           ...          ...           ...
1772063061_D09           13         162.0           17            11
1772067059_B04           13         203.5           17            11
1772066097_D04           13         266.5           17            11
1772063068_D01           13         170.0           17            11
1772066098_A12           15         159.5           19            10
```

Let's compare clusters from both graphs with each other and with cell type annotations provided by the authors, using the adjusted Rand index (ARI).

```r
# Compare clusterings with ARI
c(
    SNN_vs_kNN = pairwiseRand(ge$snn__cluster, ge$knn__cluster, mode = "index"),
    SNN_vs_annotation = pairwiseRand(ge$snn__cluster, ge$level1class, mode =
     ↪  "index"),
    kNN_vs_annotation = pairwiseRand(ge$knn__cluster, ge$level1class, mode =
     ↪  "index")
)
```

```
       SNN_vs_kNN SNN_vs_annotation kNN_vs_annotation
        0.7052230         0.4447928         0.3857959
```

```r
# Cross-tabulation of SNN clusters and annotated cell types
table(ge$snn__cluster, ge$level1class)
```

```
     astrocytes_ependymal endothelial-mural interneurons microglia
  1                     3                 4          275         0
  10                   10                 0            0         0
  11                    0                 0            0         1
  12                    0                 0            0         0
  13                   24               166            0         5
  14                    0                 1            0        79
  15                  130                 3            0         1
  2                     0                 0            7         0
  3                     0                 2            5         2
  4                     0                 0            2         0
  5                     0                 0            1         0
  6                     2                 0            0         0
  7                    17                 0            0         0
  8                     0                 1            0         0
```

```
9                      1                 0            0         1

   oligodendrocytes pyramidal CA1 pyramidal SS
1                 0             4            0
10              233             0            0
11              157             0            0
12              219             0            0
13                5             0            0
14                0             0            0
15                0             0            0
2               164             0           33
3                 1             0          199
4                 0           503            4
5                 0           199            1
6                 0            20          150
7                 0            56           10
8                 0           110            0
9                17            46            0
```

Finally, let's visualize the SNN graph using the t-SNE coordinates (stored in `reducedDims`) as layout, with cells colored by SNN clusters (a node attribute of the graph) and by annotated cell types (a variable in `colData`). Since all data are in the same object, and `colGraph()` adds variables in `colData` as node attributes, plotting is straightforward.

```
# Extract SNN graph with `colData` variables as node attributes
g <- colGraph(ge, "snn")
tsne <- reducedDim(ge, "TSNE")[V(g)$name, ]

# For visualization, keep only the strongest edges
g_plot <- delete_edges(g, which(E(g)$weight < quantile(E(g)$weight, 0.9)))

p1 <- ggraph(g_plot, layout = "manual", x = tsne[, 1], y = tsne[, 2]) +
    geom_edge_link(alpha = 0.05, color = "grey50") +
    geom_node_point(aes(color = factor(as.numeric(cluster))), size = 0.5) +
    guides(color = guide_legend(override.aes = list(size = 3))) +
    theme_graph(base_family = "sans") +
    labs(color = "SNN cluster", title = "SNN graph - Louvain clusters")

p2 <- ggraph(g_plot, layout = "manual", x = tsne[, 1], y = tsne[, 2]) +
    geom_edge_link(alpha = 0.05, color = "grey50") +
    geom_node_point(aes(color = level1class), size = 0.5) +
    scale_color_brewer(palette = "Dark2") +
    guides(color = guide_legend(override.aes = list(size = 3))) +
    theme_graph(base_family = "sans") +
    labs(color = "Cell type", title = "SNN graph - annotated cell types")

wrap_plots(p1, p2, nrow = 1) +
    plot_annotation(tag_levels = "A") &
    theme(plot.title = element_text(size = 13))
```

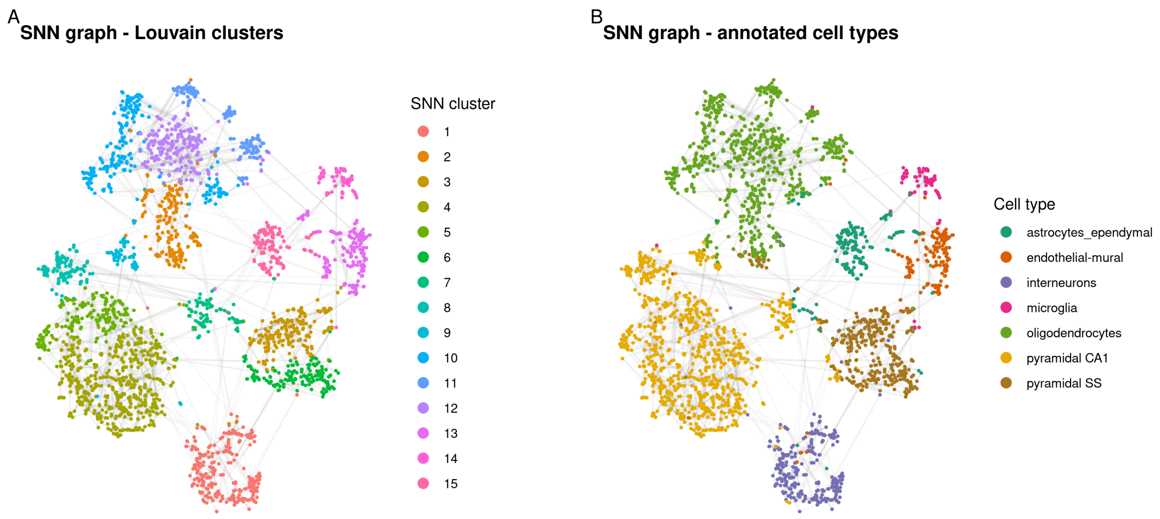

#### Subsetting

A key advantage of storing graphs in a `GraphExperiment` object is that subsetting operations are propagated to graphs. For instance, a common step in scRNA-seq data analysis is to subset a particular cell population and re-cluster it to identify subpopulations. Here, we will subset interneurons. The graphs in `colGraphs` are subset accordingly (i.e., induced subgraphs), and cluster assignments from the full graph are kept, so we can compare them with clusters from the subgraph.

```
# Subset interneurons
ge_int <- ge[, ge$level1class == "interneurons"]
ge_int
```

```
class: GraphExperiment
dim: 20006 290
metadata(2): qc PCA
assays(2): counts logcounts
rownames(20006): Tspan12 Tshz1 ... mt-Rnr1 mt-Nd4l
rowData names(6): featureType means ... residuals hvg
colnames(290): 1772071015_C02 1772071017_G12 ... 1772071014_F12
  1772071014_E02
colData names(18): tissue group # ... knn__cluster knn__strength
reducedDimNames(2): PCA TSNE
mainExpName: gene
altExpNames(2): repeat ERCC
rowGraphs(0):
colGraphs(2): snn knn
```

```
# Graphs were subset as well
sapply(colGraphs(ge_int), vcount)
```

```
snn knn
290 290
```

```r
# Re-cluster interneurons using the induced subgraph
g_int <- colGraph(ge_int, "snn")
V(g_int)$subcluster <- as.character(membership(cluster_louvain(g_int)))
colGraph(ge_int, "snn") <- g_int

# Subclusters vs annotated subtypes
table(ge_int$snn__subcluster, ge_int$level2class)
```

```
   Int1 Int10 Int11 Int12 Int13 Int14 Int15 Int16 Int2 Int3 Int4 Int5 Int6
 1    0     9     7    21     0     6     1     7    0    0    1    2    1
 2    0    10     1     0     0     0     0     0    0    1    2   13   17
 3   12     1     2     0     7     4     9     1   24    8   11    0    0
 4    0     0     0     0     8    12     8    12    0    0    0    0    0
 5    0     1     0     0     0     0     0     0    0    1    1    5    4

   Int7 Int8 Int9
 1    3    0    6
 2   16    7    2
 3    1    0    3
 4    0    0    0
 5    3   19    0
```

In summary, **GraphExperiment** makes cell-cell graphs first-class citizens of single-cell data analysis, allowing users to keep the graphs used for clustering (or for other tasks, such as trajectory inference or label propagation), along with the expression data and dimensionality reduction results, in a single object.

#### Session info

This document was created under the following conditions:

```
─ Session info ───────────────────────────────────────────────────────────────
 setting  value
 version  R version 4.5.2 (2025-10-31)
 os       Ubuntu 22.04.5 LTS
 system   x86_64, linux-gnu
 ui       X11
 language (EN)
 collate  en_US.UTF-8
 ctype    en_US.UTF-8
 tz       Europe/Brussels
 date     2026-10-24
 pandoc   3.6.3 @ /opt/quarto/bin/tools/ (via rmarkdown)
 quarto   1.8.25 @ /usr/local/bin/quarto

─ Packages ───────────────────────────────────────────────────────────────────
 package              * version date (UTC) lib source
 Biobase              * 2.70.0  2025-10-29 [1] Bioconductor 3.22 (R 4.5.2)
 BiocGenerics         * 0.56.0  2025-10-29 [1] Bioconductor 3.22 (R 4.5.2)
 bluster              * 1.20.0  2025-10-29 [1] Bioconductor 3.22 (R 4.5.2)
 dplyr                * 1.1.4   2023-11-17 [1] CRAN (R 4.5.2)
 forcats              * 1.0.1   2025-09-25 [1] CRAN (R 4.5.2)
 generics             * 0.1.4   2025-05-09 [1] CRAN (R 4.5.2)
 GenomicRanges        * 1.62.1  2025-12-08 [1] Bioconductor 3.22 (R 4.5.2)
 ggplot2              * 4.0.1   2025-11-14 [1] CRAN (R 4.5.2)
 ggraph               * 2.2.2   2025-08-24 [1] CRAN (R 4.5.2)
 GraphExperiment      * 1.1.2   2026-09-08 [1] Bioconductor
```

```
here                  * 1.0.2   2025-09-15 [1] CRAN (R 4.5.2)
igraph                * 2.2.1   2025-10-27 [1] CRAN (R 4.5.2)
IRanges               * 2.44.0  2025-10-29 [1] Bioconductor 3.22 (R 4.5.2)
lubridate             * 1.9.4   2024-12-08 [1] CRAN (R 4.5.2)
MatrixGenerics        * 1.22.0  2025-10-29 [1] Bioconductor 3.22 (R 4.5.2)
matrixStats           * 1.5.0   2025-01-07 [1] CRAN (R 4.5.2)
patchwork             * 1.3.2   2025-08-25 [1] CRAN (R 4.5.2)
purrr                 * 1.2.1   2026-01-09 [1] CRAN (R 4.5.2)
readr                 * 2.1.6   2025-11-14 [1] CRAN (R 4.5.2)
S4Vectors             * 0.48.0  2025-10-29 [1] Bioconductor 3.22 (R 4.5.2)
scrapper              * 1.7.9   2026-10-23 [1] Github (libscran/scrapper@224ce0f)
scRNAseq              * 2.24.0  2025-11-04 [1] Bioconductor 3.22 (R 4.5.2)
Seqinfo               * 1.0.0   2025-10-29 [1] Bioconductor 3.22 (R 4.5.2)
SingleCellExperiment * 1.32.0  2025-10-29 [1] Bioconductor 3.22 (R 4.5.2)
stringr               * 1.6.0   2025-11-04 [1] CRAN (R 4.5.2)
SummarizedExperiment * 1.40.0  2025-10-29 [1] Bioconductor 3.22 (R 4.5.2)
tibble                * 3.3.1   2026-01-11 [1] CRAN (R 4.5.2)
tidyr                 * 1.3.2   2025-12-19 [1] CRAN (R 4.5.2)
tidyverse             * 2.0.0   2023-02-22 [1] CRAN (R 4.5.2)

[1] /home/faalm/R/x86_64-pc-linux-gnu-library/4.5
[2] /usr/local/lib/R/site-library
[3] /usr/lib/R/site-library
[4] /usr/lib/R/library
* -- Packages attached to the search path.

------------------------------------------------------------------------------
```

### Use case 5: cell-cell communication networks

Cell-cell communication (CCC) is commonly inferred from scRNA-seq data based on the expression of ligands in sender cell types and their receptors in receiver cell types. The result is naturally represented as a directed graph of cell types. Here, we will use **GraphExperiment** to store CCC networks in the `colGraphs` slot of an object containing pseudobulk expression profiles (i.e., with cell types in columns). This use case demonstrates that observation-level graphs can be stored at any resolution, including cells (see use case 4), cell types, samples, or species (see use cases 6 and 7).

We will infer CCC with **LIANA** (Dimitrov et al. 2022), a framework that runs multiple CCC inference methods and combines their results into a consensus ranking of ligand-receptor interactions. We chose LIANA because a systematic comparison of CCC methods showed that no single method consistently outperformed the others, and that the choice of method strongly affects predictions (Dimitrov et al. 2022), making a consensus approach a sensible default. Other tools, such as NicheNet (Browaeys, Saelens, and Saeys 2019), address a complementary question (i.e., which ligands best explain gene expression changes in receiver cells), and their output (ligand-target gene links) could be stored in `rowGraphs`. We will use scRNA-seq data of human pancreatic islets from Baron et al. (2016), available in the **scRNAseq** package (Risso and Cole 2025).

```r
library(here)
library(GraphExperiment)
library(scRNAseq)
library(scuttle)
library(liana)
library(tidyverse)
library(ggraph)

set.seed(123) # for reproducibility
```

#### Data description and processing

We will first load the data and, to keep this example lightweight, randomly sample up to 100 cells per cell type, keeping only cell types with at least 20 cells.

```r
# Load data
sce <- BaronPancreasData("human")
sce
```

```
class: SingleCellExperiment
dim: 20125 8569
metadata(0):
assays(1): counts
rownames(20125): A1BG A1CF ... ZZZ3 pk
rowData names(0):
colnames(8569): human1_lib1.final_cell_0001 human1_lib1.final_cell_0002
  ... human4_lib3.final_cell_0700 human4_lib3.final_cell_0701
colData names(2): donor label

reducedDimNames(0):
mainExpName: NULL
```

```
altExpNames(0):
```

```
# Check annotated cell types
table(sce$label)
```

```
           acinar activated_stellate              alpha               beta 
              958                284               2326               2525 
            delta             ductal        endothelial            epsilon 
              601               1077                252                 18 
            gamma         macrophage               mast quiescent_stellate 
              255                 55                 25                173 
          schwann             t_cell 
               13                  7 
```

```
# Sample up to 100 cells per cell type
idx <- split(seq_len(ncol(sce)), sce$label)
idx <- unlist(lapply(idx, function(i) if(length(i) > 100) sample(i, 100) else i))
sce <- sce[, sort(idx)]

# Keep only cell types with at least 20 cells
sce <- sce[, sce$label %in% names(which(table(sce$label) >= 20))]
table(sce$label)
```

```
           acinar activated_stellate              alpha               beta 
              100                100                100                100 
            delta             ductal        endothelial              gamma 
              100                100                100                100 
       macrophage               mast quiescent_stellate 
               55                 25                100 
```

```
# Normalize data
sce <- logNormCounts(sce)

sce
```

```
class: SingleCellExperiment 
dim: 20125 980 
metadata(0):
assays(2): counts logcounts
rownames(20125): A1BG A1CF ... ZZZ3 pk
rowData names(0):
colnames(980): human1_lib1.final_cell_0016 human1_lib1.final_cell_0028
  ... human4_lib3.final_cell_0683 human4_lib3.final_cell_0697
colData names(3): donor label sizeFactor

reducedDimNames(0):
mainExpName: NULL

altExpNames(0):
```

#### Inferring cell-cell communication

To reduce memory usage, we will only keep genes encoding ligands and receptors in LIANA's consensus resource. Then, we will infer CCC with four methods (NATMI, Connectome, log2FC, and SingleCellSignalR), and aggregate their results into a consensus ranking.

```r
# Keep only genes in the ligand-receptor resource
lr_resource <- select_resource("Consensus")[["Consensus"]]
lr_genes <- unique(unlist(strsplit(
    c(lr_resource$source_genesymbol, lr_resource$target_genesymbol), "_"
)))
sce_lr <- sce[rownames(sce) %in% lr_genes, ]

# Infer CCC with multiple methods and aggregate results
liana_res <- liana_wrap(
    sce_lr, idents_col = "label", resource = "Consensus",
    method = c("natmi", "connectome", "logfc", "sca"), verbose = FALSE
)
liana_agg <- liana_aggregate(liana_res, verbose = FALSE)

head(liana_agg)
```

```
# A tibble: 6 x 14
  source         target ligand.complex receptor.complex aggregate_rank mean_rank
  <chr>          <chr>  <chr>          <chr>                     <dbl>     <dbl>
1 activated_ste~ endot~ COL1A2         CD93                   7.59e-10      50.8
2 activated_ste~ endot~ COL1A1         CD93                   9.72e-10      38.2
3 alpha          mast   GCG            ADRB2                  1.46e- 8      74.8
4 endothelial    endot~ COL4A1         CD93                   3.39e- 8     168
5 activated_ste~ endot~ MMP2           PECAM1                 3.74e- 8     119
6 alpha          endot~ GCG            RAMP2                  4.11e- 8      94.5
# i 8 more variables: natmi.edge_specificity <dbl>, natmi.rank <dbl>,
#   connectome.weight_sc <dbl>, connectome.rank <dbl>, logfc.logfc_comb <dbl>,
#   logfc.rank <dbl>, sca.LRscore <dbl>, sca.rank <dbl>
```

We will keep interactions with aggregate rank $< 0.01$ and create a directed graph of cell types in which edges point from sender to receiver cell types, edge weights represent the number of interactions, and edge attributes include the ligand-receptor pairs. Self-loops represent autocrine signaling.

```r
# Filter interactions and summarize by sender/receiver pairs
ccc_edges <- liana_agg |>
    filter(aggregate_rank < 0.01) |>
    mutate(lr_pair = paste0(ligand.complex, "->", receptor.complex)) |>
    group_by(source, target) |>
    summarise(
        weight = n(),
        lr_pairs = paste(lr_pair, collapse = ";"),
        .groups = "drop"
    )

head(ccc_edges)
```

```
# A tibble: 6 x 4
  source target             weight lr_pairs
  <chr>  <chr>               <int> <chr>
1 acinar acinar                  9 PRSS2->PARD3;PRSS1->PARD3;PRSS3->PARD3;LAMB3~
2 acinar activated_stellate     16 PRSS2->F2R;PRSS1->F2R;PRSS3->F2R;PRSS1->F2RL~
3 acinar beta                    3 PRSS2->PARD3;PRSS1->PARD3;PRSS3->PARD3
4 acinar ductal                 17 PRSS2->PARD3;SPINK1->EGFR;PRSS1->F2RL1;PRSS3~
5 acinar endothelial            16 PRSS2->F2R;CEL->CXCR4;PRSS1->F2R;ANXA2->ROBO~
6 acinar macrophage              7 ALB->B2M_FCGRT;LCN2->FCGR2A;PLTP->ABCA1;RPS1~
```

#### Storing pseudobulk data and CCC networks in a `GraphExperiment` object

Next, we will aggregate counts by cell type to obtain pseudobulk profiles, so that observations (columns) are cell types. Then, we will create the CCC graph, including all cell types as nodes, and add the total number of outgoing and incoming interactions (i.e., the weighted out- and in-degrees) as node attributes, which indicate the roles of cell types as senders and receivers, respectively.

```
# Create pseudobulk profiles
pb <- aggregateAcrossCells(
    sce, ids = sce$label, statistics = "sum", use.assay.type = "counts"
)
sizeFactors(pb) <- NULL # remove cell-level size factors
pb <- logNormCounts(pb)

pb
```

```
class: SingleCellExperiment
dim: 20125 11
metadata(0):
assays(2): counts logcounts
rownames(20125): A1BG A1CF ... ZZZ3 pk
rowData names(0):
colnames(11): acinar activated_stellate ... mast quiescent_stellate
colData names(5): donor label ids ncells sizeFactor

reducedDimNames(0):
mainExpName: NULL

altExpNames(0):
```

```
# Create CCC graph with all cell types as nodes
ccc_graph <- graph_from_data_frame(
    ccc_edges, directed = TRUE, vertices = data.frame(name = colnames(pb))
)
V(ccc_graph)$out_strength <- strength(ccc_graph, mode = "out")
V(ccc_graph)$in_strength <- strength(ccc_graph, mode = "in")

# Create `GraphExperiment` object
ge <- as(pb, "GraphExperiment")
colGraph(ge, "ccc") <- ccc_graph

ge
```

```
class: GraphExperiment
dim: 20125 11
metadata(0):
assays(2): counts logcounts
rownames(20125): A1BG A1CF ... ZZZ3 pk
rowData names(0):
colnames(11): acinar activated_stellate ... mast quiescent_stellate
colData names(7): donor label ... ccc__out_strength ccc__in_strength

reducedDimNames(0):
mainExpName: NULL
```

```
altExpNames(0):
rowGraphs(0):
colGraphs(1): ccc
```

Node attributes are available in `colData`, along with cell type metadata (e.g., number of cells per cell type).

```
colData(ge) |>
    as.data.frame() |>
    select(ncells, ccc__out_strength, ccc__in_strength) |>
    arrange(-ccc__out_strength)
```

```
                   ncells ccc__out_strength ccc__in_strength
activated_stellate    100               755              389
quiescent_stellate    100               347              235
endothelial           100               299              496
ductal                100               259              442
macrophage             55               242              303
beta                  100                93               44
alpha                 100                88               75
mast                   25                86              140
acinar                100                72              145
gamma                 100                66               55
delta                 100                58               41
```

#### Exploring the CCC network

Let's visualize the CCC network, with edge widths proportional to the number of interactions, and node sizes proportional to the number of outgoing interactions.

```
g <- colGraph(ge, "ccc")

ggraph(g, layout = "circle") +
    geom_edge_fan(
        aes(width = weight, alpha = weight),
        arrow = arrow(length = unit(2.5, "mm"), type = "closed"),
        end_cap = circle(6, "mm"), color = "steelblue"
    ) +
    geom_edge_loop(aes(width = weight, alpha = weight), color = "steelblue") +
    geom_node_point(aes(size = out_strength), color = "firebrick") +
    geom_node_text(aes(label = name), repel = TRUE, size = 4) +
    scale_edge_width(range = c(0.3, 2.5)) +
    scale_edge_alpha(range = c(0.3, 0.9), guide = "none") +
    scale_size_continuous(range = c(2, 10)) +
    theme_graph(base_family = "sans") +
    labs(edge_width = "Number of\ninteractions", size = "Outgoing\ninteractions")
```

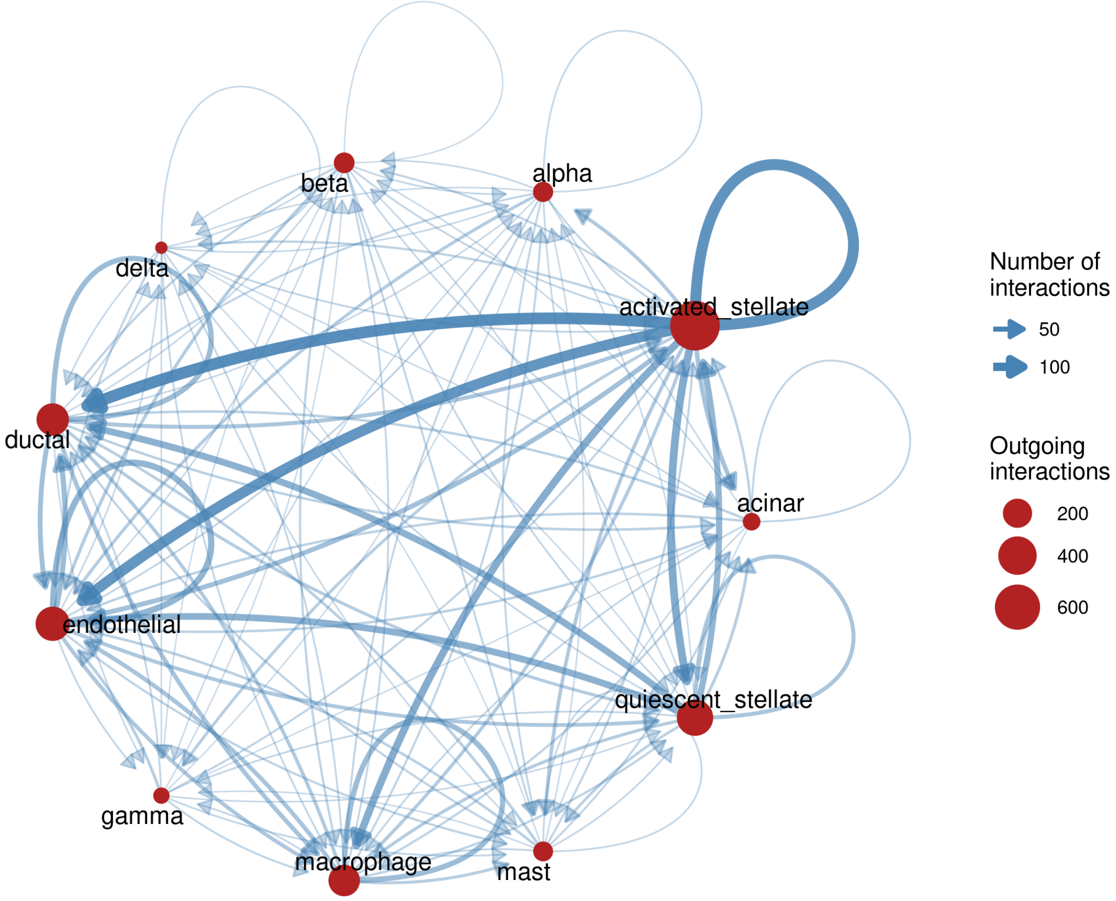

Since ligand-receptor pairs are stored as edge attributes, we can easily explore which interactions underlie each edge. For instance, let's see the ligand-receptor pairs between activated stellate cells (the main senders) and endothelial cells.

```
# Get ligand-receptor pairs for a particular edge
e <- E(g)[.from("activated_stellate") & .to("endothelial")]
lr_pairs <- strsplit(e$lr_pairs, ";")[[1]]
length(lr_pairs)
```

```
[1] 133
```

```
head(lr_pairs, 15)
```

```
 [1] "COL1A2->CD93"    "COL1A1->CD93"    "MMP2->PECAM1"    "COL18A1->KDR"
 [5] "COL1A1->ITGA5"   "SPON2->ITGA5"    "COL4A1->CD93"    "TIMP3->KDR"
 [9] "FN1->ROBO4"      "INHBA->ENG"      "TGFB3->ENG"      "VEGFA->FLT1_KDR"
[13] "COL1A1->FLT4"    "COL1A2->FLT4"    "SPARC->ENG"
```

Graph algorithms can also be used to explore the network. For instance, we can use PageRank to identify the most important receivers of signals, and then check the expression of the receptors involved in the pseudobulk profiles, which are stored in the same object.

```r
# PageRank centrality (receivers of many signals from important senders)
sort(page_rank(g, weights = E(g)$weight)$vector, decreasing = TRUE)
```

```
       endothelial             ductal activated_stellate         macrophage 
        0.19669204         0.17254930         0.14852176         0.12900457 
quiescent_stellate             acinar               mast              alpha 
        0.09092203         0.06861486         0.06562312         0.03801204 
             gamma               beta              delta 
        0.03148288         0.03052871         0.02804868 
```

```r
# Expression of receptors targeted by activated stellate cells
receptors <- liana_agg |>
    filter(aggregate_rank < 0.01, source == "activated_stellate") |>
    pull(receptor.complex) |>
    strsplit("_") |>
    unlist() |>
    unique()

round(logcounts(ge)[intersect(receptors, rownames(ge)), ], 1) |> head(10)
```

```
       acinar activated_stellate alpha beta delta ductal endothelial gamma
CD93      0.0                3.7   1.0  0.0   0.0    0.0         9.3   0.0
PECAM1    0.0                2.2   0.0  0.9   0.0    0.0        10.0   0.0
KDR       0.0                0.0   0.0  0.0   0.0    0.0         9.0   0.0
ITGA5     2.0                7.9   0.0  0.0   0.0    6.4         9.4   1.1
CD44      9.9                7.2   5.7  4.8   3.9    9.4         4.7   5.5
SDC1      2.6                1.7   0.0  0.0   0.0    5.9         0.0   0.0
ROBO4     0.0                0.0   0.0  0.0   0.0    0.0         7.9   0.0
DDR2      1.6                6.7   0.0  0.0   2.7    0.0         2.8   0.0
ENG       0.9                8.3   0.0  0.9   0.0    1.4        11.1   1.1
ITGB2     0.0                0.0   0.0  0.0   0.0    0.9         1.3   0.0
       macrophage mast quiescent_stellate
CD93          4.7  0.0                3.5
PECAM1        7.0  0.0                1.3
KDR           0.0  0.0                0.0
ITGA5         6.3  5.4                6.3
CD44          7.5  8.9                6.5
SDC1          0.0  0.0                1.3
ROBO4         0.0  0.0                0.0
DDR2          0.0  0.0                5.9
ENG           8.6  7.1                8.6
ITGB2         8.6  3.5                0.0
```

#### Session info

This document was created under the following conditions:

```
- Session info ---------------------------------------------------------------
 setting  value
 version  R version 4.5.2 (2025-10-31)
 os       Ubuntu 22.04.5 LTS
 system   x86_64, linux-gnu
 ui       X11
 language (EN)
 collate  en_US.UTF-8
```

```
 ctype    en_US.UTF-8
 tz       Europe/Brussels
 date     2026-10-24
 pandoc   3.6.3 @ /opt/quarto/bin/tools/ (via rmarkdown)
 quarto   1.8.25 @ /usr/local/bin/quarto

- Packages -------------------------------------------------------------------
 package              * version date (UTC) lib source
 Biobase              * 2.70.0  2025-10-29 [1] Bioconductor 3.22 (R 4.5.2)
 BiocGenerics         * 0.56.0  2025-10-29 [1] Bioconductor 3.22 (R 4.5.2)
 dplyr                * 1.1.4   2023-11-17 [1] CRAN (R 4.5.2)
 forcats              * 1.0.1   2025-09-25 [1] CRAN (R 4.5.2)
 generics             * 0.1.4   2025-05-09 [1] CRAN (R 4.5.2)
 GenomicRanges        * 1.62.1  2025-12-08 [1] Bioconductor 3.22 (R 4.5.2)
 ggplot2              * 4.0.1   2025-11-14 [1] CRAN (R 4.5.2)
 ggraph               * 2.2.2   2025-08-24 [1] CRAN (R 4.5.2)
 GraphExperiment      * 1.1.2   2026-09-08 [1] Bioconductor
 here                 * 1.0.2   2025-09-15 [1] CRAN (R 4.5.2)
 igraph               * 2.2.1   2025-10-27 [1] CRAN (R 4.5.2)
 IRanges              * 2.44.0  2025-10-29 [1] Bioconductor 3.22 (R 4.5.2)
 liana                * 0.1.14  2026-10-21 [1] Github (saezlab/liana@6cab46c)
 lubridate            * 1.9.4   2024-12-08 [1] CRAN (R 4.5.2)
 MatrixGenerics       * 1.22.0  2025-10-29 [1] Bioconductor 3.22 (R 4.5.2)
 matrixStats          * 1.5.0   2025-01-07 [1] CRAN (R 4.5.2)
 purrr                * 1.2.1   2026-01-09 [1] CRAN (R 4.5.2)
 readr                * 2.1.6   2025-11-14 [1] CRAN (R 4.5.2)
 S4Vectors            * 0.48.0  2025-10-29 [1] Bioconductor 3.22 (R 4.5.2)
 scRNAseq             * 2.24.0  2025-11-04 [1] Bioconductor 3.22 (R 4.5.2)
 scuttle              * 1.20.0  2025-10-30 [1] Bioconductor 3.22 (R 4.5.2)
 Seqinfo              * 1.0.0   2025-10-29 [1] Bioconductor 3.22 (R 4.5.2)
 SingleCellExperiment * 1.32.0  2025-10-29 [1] Bioconductor 3.22 (R 4.5.2)
 stringr              * 1.6.0   2025-11-04 [1] CRAN (R 4.5.2)
 SummarizedExperiment * 1.40.0  2025-10-29 [1] Bioconductor 3.22 (R 4.5.2)
 tibble               * 3.3.1   2026-01-11 [1] CRAN (R 4.5.2)
 tidyr                * 1.3.2   2025-12-19 [1] CRAN (R 4.5.2)
 tidyverse            * 2.0.0   2023-02-22 [1] CRAN (R 4.5.2)

 [1] /home/faalm/R/x86_64-pc-linux-gnu-library/4.5
 [2] /usr/local/lib/R/site-library
 [3] /usr/lib/R/site-library
 [4] /usr/lib/R/library
 * -- Packages attached to the search path.

------------------------------------------------------------------------------
```

### Use case 6: comparative genomics with species and phylogenetic profile graphs

In comparative genomics, data are often represented as matrices with features (e.g., gene families, orthogroups, synteny clusters) in rows and species in columns. Here, we will use **GraphExperiment** to store a matrix of *phylogenomic profiles* (i.e., copy number of synteny clusters across species) along with two graphs: a species-species graph derived from a phylogeny (in `colGraphs`), and a graph of synteny clusters with similar phylogenetic profiles (in `rowGraphs`). We will use data from the **syntenet** package (Almeida-Silva et al. 2022), which include synteny network clusters for 25 eudicot species from Zhao and Schranz (2019), and a microsynteny-based angiosperm phylogeny (Zhao et al. 2021).

We will also demonstrate how phylogenies (`phylo` objects) can be represented as `igraph` objects, since phylogenies are a special kind of graph (i.e., a directed acyclic graph, DAG).

```
library(here)
library(GraphExperiment)
library(syntenet)
library(tidyverse)
library(ggraph)
library(ComplexHeatmap)
library(ape)

set.seed(123) # for reproducibility
```

#### Data description

The data set `clusters` is a data frame with genes and the synteny clusters they belong to, and the species tree `angiosperm_phylogeny` is a `phylo` object with 123 angiosperm species. We will first create a phylogenomic profiles matrix with the copy number of each synteny cluster across species.

```
# Load data
data(clusters)
data(angiosperm_phylogeny)

head(clusters)
```

```
       Gene Cluster
1 cca_23646      25
2 cca_23668    1089
3 cca_32926     304
4 cca_26186     254
5 cca_24381     592
6 cca_24396      20
```

```
angiosperm_phylogeny
```

```
Phylogenetic tree with 123 tips and 121 internal nodes.
```

```
Tip labels:
  Prunus mume, Prunus persica, Pyrus x bretschneideri, Malus domestica, Rosa chinensis, Fragaria vesca
Node labels:
  , 100/100, 100/100, 100/100, 100/100, 100/100, ...

Unrooted; includes branch length(s).
```

```r
# Get phylogenomic profiles
profiles <- phylogenomic_profile(clusters)
profiles <- matrix(profiles, nrow = nrow(profiles), dimnames = dimnames(profiles))
rownames(profiles) <- paste0("SC", rownames(profiles))
profiles[1:5, 1:5]
```

```
    adu car cca fve gma
SC1   0   1   2   2   3
SC2   2   2   2   1   3
SC3   1   1   1   1   2
SC4   1   2   1   1   4
SC5   1   1   1   1   1
```

Species in the profiles matrix are represented by abbreviations, while species in the tree are represented by their full names. Thus, we will create a data frame with species metadata (full names and families), and keep only species that are present in both data sets.

```r
# Create species metadata
species_md <- tribble(
    ~abbrev, ~species, ~family,
    "adu", "Arachis duranensis", "Fabaceae",
    "car", "Cicer arietinum", "Fabaceae",
    "cca", "Cercis canadensis", "Fabaceae",
    "fve", "Fragaria vesca", "Rosaceae",
    "gma", "Glycine max", "Fabaceae",
    "hlu", "Humulus lupulus", "Cannabaceae",
    "jcu", "Jatropha curcas", "Euphorbiaceae",
    "Lang", "Lupinus angustifolius", "Fabaceae",
    "lja", "Lotus japonicus", "Fabaceae",
    "lus", "Linum usitatissimum", "Linaceae",
    "mdo", "Malus domestica", "Rosaceae",
    "mes", "Manihot esculenta", "Euphorbiaceae",
    "Mnot", "Morus notabilis", "Moraceae",
    "mtr", "Medicago truncatula", "Fabaceae",
    "pbr", "Pyrus x bretschneideri", "Rosaceae",
    "pmu", "Prunus mume", "Rosaceae",
    "ppe", "Prunus persica", "Rosaceae",
    "ptr", "Populus trichocarpa", "Salicaceae",
    "pvu", "Phaseolus vulgaris", "Fabaceae",
    "rco", "Ricinus communis", "Euphorbiaceae",
    "tpr", "Trifolium pratense", "Fabaceae",
    "van", "Vigna angularis", "Fabaceae",
    "vra", "Vigna radiata", "Fabaceae",
    "Zjuj", "Ziziphus jujuba", "Rhamnaceae"
) |>
    filter(species %in% angiosperm_phylogeny$tip.label) |>
    column_to_rownames("species")

nrow(species_md)
```

```
[1] 21
```

```r
# Keep only species in both data sets and rename columns to full names
profiles <- profiles[, species_md$abbrev]
colnames(profiles) <- rownames(species_md)

# Remove synteny clusters that are absent from all remaining species
profiles <- profiles[rowSums(profiles) > 0, ]

# Quick look at the final profile matrix
dim(profiles)
```

```
[1] 2535   21
```

```r
profiles[1:5, 1:5]
```

```
    Arachis duranensis Cicer arietinum Fragaria vesca Glycine max
SC1                  0               1              2           3
SC2                  2               2              1           3
SC3                  1               1              1           2
SC4                  1               2              1           4
SC5                  1               1              1           1

    Lupinus angustifolius
SC1                     2
SC2                     3
SC3                     1
SC4                     2
SC5                     2
```

#### Phylogenies as graphs

A phylogeny is a DAG in which edges point from ancestors to descendants. Thus, `phylo` objects can beconverted to `igraph` objects with the `as.igraph()` method from **ape** (Paradis and Schliep 2018). Here, we will first subset the tree to keep only our species of interest, and then convert it to a DAG with branch lengths as edge attributes.

```r
# Subset tree and remove node labels (bootstrap support values)
tree <- ape::keep.tip(angiosperm_phylogeny, colnames(profiles))
tree$node.label <- NULL

# Convert `phylo` to `igraph`, adding branch lengths as edge attributes
dag <- ape::as.igraph.phylo(tree, directed = TRUE)
E(dag)$length <- tree$edge.length
V(dag)$is_tip <- V(dag)$name %in% tree$tip.label

dag
```

```
IGRAPH f4f1712 DN-- 41 40 --
+ attr: name (v/c), is_tip (v/l), length (e/n)
+ edges from f4f1712 (vertex names):
 [1] Node1 ->Node2                  Node2 ->Node3
 [3] Node3 ->Node4                  Node4 ->Node5
 [5] Node5 ->Node6                  Node6 ->Prunus mume
 [7] Node6 ->Prunus persica         Node5 ->Node7
 [9] Node7 ->Pyrus x bretschneideri Node7 ->Malus domestica
```

```
[11] Node4 ->Fragaria vesca         Node3 ->Node8
[13] Node8 ->Morus notabilis        Node8 ->Ziziphus jujuba
[15] Node2 ->Node9                  Node9 ->Node10
+ ... omitted several edges
```

```r
is_dag(dag)
```

```
[1] TRUE
```

Since the tree is now an `igraph` object, any graph algorithm can be used on it. For instance, we can obtain the matrix of shared evolutionary history between pairs of species (i.e., the distance from the root to their most recent common ancestor), which is the variance-covariance matrix under a Brownian motion model used by most phylogenetic comparative methods. For tips $i$ and $j$, the shared path length is $(d_{root,i} + d_{root,j} - d_{i,j})/2$, where $d$ is the distance between nodes along branches.

```r
# Get distances between all pairs of nodes
d <- distances(dag, mode = "all", weights = E(dag)$length)
root <- V(dag)[igraph::degree(dag, mode = "in") == 0]$name
tips <- V(dag)$name[V(dag)$is_tip]

# Get shared path lengths
shared <- outer(d[root, tips], d[root, tips], "+") / 2 - d[tips, tips] / 2

# Sanity check: same as the output of `ape::vcv()`
all.equal(shared, ape::vcv(tree)[tips, tips])
```

```
[1] TRUE
```

Nodes of graphs in `colGraphs` must match column names (here, species) *exactly*. Thus, DAGs representing phylogenies (which include internal nodes) cannot be stored in the `colGraphs` slot. Instead, we will store a species-species graph with edge weights representing shared evolutionary history (i.e., the off-diagonal elements of the matrix above), and a node attribute (`root_distance`) with the distances from the root to each species (i.e., the diagonal). Pairs of species whose most recent common ancestor is the root share no evolutionary history (weight of zero), so they are not connected.

> **Note**
>
> Native support for graphs containing nodes that are not observations (e.g., internal nodes of phylogenies and phylogenetic networks) is planned for future versions of **GraphExperiment**. For now, full DAGs can be stored in the `metadata` slot, as we will do below.

```r
# Create species graph from the matrix of shared path lengths
phylo_graph <- graph_from_adjacency_matrix(
    shared, mode = "undirected", weighted = TRUE, diag = FALSE
)
V(phylo_graph)$root_distance <- diag(shared)[V(phylo_graph)$name]

phylo_graph
```

```
IGRAPH 3361dc9 UNW- 21 174 --
+ attr: name (v/c), root_distance (v/n), weight (e/n)
+ edges from 3361dc9 (vertex names):
 [1] Prunus mume--Prunus persica         Prunus mume--Pyrus x bretschneideri
 [3] Prunus mume--Malus domestica        Prunus mume--Fragaria vesca
 [5] Prunus mume--Morus notabilis        Prunus mume--Ziziphus jujuba
```

```
 [7] Prunus mume--Vigna radiata         Prunus mume--Vigna angularis
 [9] Prunus mume--Phaseolus vulgaris    Prunus mume--Glycine max
[11] Prunus mume--Trifolium pratense    Prunus mume--Medicago truncatula
[13] Prunus mume--Cicer arietinum       Prunus mume--Lotus japonicus
[15] Prunus mume--Lupinus angustifolius  Prunus mume--Arachis duranensis
+ ... omitted several edges
```

#### Phylogenetic profile similarity graph

Synteny clusters (or gene families) with similar patterns of presence and absence across species (i.e., similar phylogenetic profiles) are likely functionally related, as genes in the same pathway or protein complex tend to be gained and lost together (Pellegrini et al. 1999). Here, we will create a graph of synteny clusters by connecting clusters with Jaccard similarities $\geq 0.9$ between their presence/absence profiles. Since clusters present in (nearly) all species or in very few species provide little information, we will only consider clusters that are absent in at least three species and present in at least three species. All other clusters will be included in the graph as isolated nodes, since nodes in row graphs must match row names *exactly*.

```
# Get presence/absence profiles of informative clusters
pa <- (profiles > 0) * 1
n_species <- ncol(pa)
informative <- rowSums(pa) >= 3 & rowSums(pa) <= n_species - 3
sum(informative)
```

```
[1] 1400
```

```
# Get Jaccard similarities between informative clusters
jaccard <- 1 - as.matrix(dist(pa[informative, ], method = "binary"))
jaccard[jaccard < 0.9] <- 0

# Create graph, including all clusters as nodes
profile_graph <- graph_from_adjacency_matrix(
    jaccard, mode = "undirected", weighted = TRUE, diag = FALSE
) |>
    add_vertices(
        sum(!informative), name = rownames(pa)[!informative]
    )

profile_graph
```

```
IGRAPH 30793ad UNW- 2535 3044 --
+ attr: name (v/c), weight (e/n)
+ edges from 30793ad (vertex names):
 [1] SC10 --SC84   SC11 --SC232  SC11 --SC382  SC11 --SC464  SC19 --SC1172
 [6] SC24 --SC26   SC24 --SC329  SC24 --SC497  SC24 --SC787  SC24 --SC1060
[11] SC24 --SC1281 SC26 --SC329  SC26 --SC497  SC26 --SC567  SC26 --SC787
[16] SC26 --SC813  SC26 --SC1001 SC26 --SC1060 SC57 --SC120  SC57 --SC658
[21] SC57 --SC683  SC61 --SC260  SC61 --SC342  SC61 --SC363  SC61 --SC465
[26] SC61 --SC1263 SC67 --SC235  SC85 --SC654  SC94 --SC496  SC95 --SC1230
[31] SC105--SC474  SC105--SC899  SC118--SC155  SC118--SC463  SC118--SC766
[36] SC118--SC972  SC120--SC658  SC120--SC683  SC121--SC180  SC121--SC268
+ ... omitted several edges
```

We will then identify modules (or communities) of synteny clusters with similar profiles using the Louvain algorithm.

```r
# Identify modules of clusters with similar profiles (Louvain algorithm)
modules <- as.character(membership(cluster_louvain(profile_graph)))
modules[igraph::degree(profile_graph) == 0] <- NA # isolated nodes are not in
 ↪  modules
V(profile_graph)$module <- modules

profile_graph
```

```
IGRAPH 30793ad UNW- 2535 3044 --
+ attr: name (v/c), module (v/c), weight (e/n)
+ edges from 30793ad (vertex names):
 [1] SC10 --SC84   SC11 --SC232  SC11 --SC382  SC11 --SC464  SC19 --SC1172
 [6] SC24 --SC26   SC24 --SC329  SC24 --SC497  SC24 --SC787  SC24 --SC1060
[11] SC24 --SC1281 SC26 --SC329  SC26 --SC497  SC26 --SC567  SC26 --SC787
[16] SC26 --SC813  SC26 --SC1001 SC26 --SC1060 SC57 --SC120  SC57 --SC658
[21] SC57 --SC683  SC61 --SC260  SC61 --SC342  SC61 --SC363  SC61 --SC465
[26] SC61 --SC1263 SC67 --SC235  SC85 --SC654  SC94 --SC496  SC95 --SC1230
[31] SC105--SC474  SC105--SC899  SC118--SC155  SC118--SC463  SC118--SC766
[36] SC118--SC972  SC120--SC658  SC120--SC683  SC121--SC180  SC121--SC268
+ ... omitted several edges
```

#### Storing all data in a `GraphExperiment` object

Now, we can store the phylogenomic profiles, the species metadata, and both graphs in a `GraphExperiment` object. The full DAG of the phylogeny will be stored in the `metadata` slot.

```r
ge <- GraphExperiment(
    assays = list(counts = profiles),
    colData = species_md[colnames(profiles), ],
    rowGraphs = list(profile_sim = profile_graph),
    colGraphs = list(phylo = phylo_graph),
    metadata = list(phylo_dag = dag)
)

ge
```

```
class: GraphExperiment
dim: 2535 21
metadata(1): phylo_dag
assays(1): counts
rownames(2535): SC1 SC2 ... SC5596 SC5597
rowData names(1): profile_sim__module
colnames(21): Arachis duranensis Cicer arietinum ... Vigna radiata
  Ziziphus jujuba
colData names(3): abbrev family phylo__root_distance
reducedDimNames(0):
mainExpName: NULL
altExpNames(0):
rowGraphs(1): profile_sim
colGraphs(1): phylo
```

As expected, node attributes from `rowGraphs` and `colGraphs` were added to `rowData` and `colData`, respectively.

```r
head(rowData(ge))
```

```
DataFrame with 6 rows and 1 column
    profile_sim__module
            <character>
SC1                  NA
SC2                  NA
SC3                  NA
SC4                  NA
SC5                  NA
SC6                  NA
```

```
head(colData(ge))
```

```
DataFrame with 6 rows and 3 columns
                           abbrev      family phylo__root_distance
                      <character> <character>            <numeric>
Arachis duranensis            adu    Fabaceae            0.0495666
Cicer arietinum               car    Fabaceae            0.0638300
Fragaria vesca                fve    Rosaceae            0.0437938
Glycine max                   gma    Fabaceae            0.0433086
Lupinus angustifolius        Lang    Fabaceae            0.0474296
Lotus japonicus               lja    Fabaceae            0.0612377
```

#### Phylogenetic signal of synteny clusters

Finally, we will use the species graph in `colGraphs` to quantify the phylogenetic signal in the copy number of each synteny cluster, i.e., to what extent closely related species have similar copy numbers. For that, we will calculate Moran's I (Gittleman and Kot 1990), using as weights the shared evolutionary history between species, which we can extract directly from the graph as a weighted adjacency matrix.

```
# Get weight matrix from the species graph
W <- as_adjacency_matrix(colGraph(ge, "phylo"), attr = "weight", sparse = FALSE)
W <- W[colnames(ge), colnames(ge)]

# Calculate Moran's I for each cluster with variable copy number
counts <- assay(ge, "counts")
variable <- apply(counts, 1, var) > 0
moran <- apply(counts[variable, ], 1, function(copies) {
    res <- ape::Moran.I(copies, W)
    c(I = res$observed, P = res$p.value)
})
moran <- as.data.frame(t(moran))
moran$padj <- p.adjust(moran$P, method = "BH")

# Add results to `rowData`
rowData(ge)$moran_I <- moran[rownames(ge), "I"]
rowData(ge)$moran_padj <- moran[rownames(ge), "padj"]
head(rowData(ge))
```

```
DataFrame with 6 rows and 5 columns
    profile_sim__module     moran_I profile_sim__module moran_padj
            <character>   <numeric>         <character>  <numeric>
SC1                  NA  0.15168836                  NA 0.18228154
SC2                  NA  0.28143940                  NA 0.00998879
SC3                  NA -0.02704232                  NA 0.93999366
SC4                  NA  0.03905267                  NA 0.59245157
SC5                  NA  0.00854976                  NA 0.76677411
```

```
SC6                    NA -0.05368877                 NA 0.98663195
    profile_sim__module
            <character>
SC1                    NA
SC2                    NA
SC3                    NA
SC4                    NA
SC5                    NA
SC6                    NA
```

```
# Number of clusters with significant phylogenetic signal
sum(rowData(ge)$moran_padj < 0.05, na.rm = TRUE)
```

```
[1] 629
```

Synteny clusters with phylogenetic signal include, for instance, clusters that are specific to (or lost in) particular lineages. Since the results of the Moran's I test and the module assignments in the phylogenetic profile graph are both in `rowData`, we can easily check which modules contain more clusters with significant phylogenetic signal.

```
# Modules with the largest number of clusters with phylogenetic signal
rowData(ge) |>
    as.data.frame() |>
    filter(moran_padj < 0.05, !is.na(profile_sim__module)) |>
    count(profile_sim__module, sort = TRUE) |>
    head()
```

```
  profile_sim__module  n
1                  20 73
2                 496 37
3                 660 13
4                 657 12
5                  15 10
6                 612 10
```

Let's visualize the copy number of synteny clusters with significant phylogenetic signal, with species ordered as in the phylogeny and clusters grouped by module in the phylogenetic profile graph.

```
# Subset object to keep only clusters with phylogenetic signal
ge_signal <- ge[which(rowData(ge)$moran_padj < 0.05), ]

# Get species order from the phylogeny (stored in `metadata`)
species_order <- intersect(
    names(dfs(metadata(ge)$phylo_dag, root, mode = "out")$order),
    colnames(ge_signal)
)

# Keep only the 5 largest modules for visualization
top_modules <- names(sort(table(rowData(ge_signal)$profile_sim__module), decreasing
 ↪  = TRUE))[1:5]
mod <- rowData(ge_signal)$profile_sim__module
mod[!mod %in% top_modules] <- "Other"
mod[mod != "Other"] <- paste0("M", mod[mod != "Other"])

# Plot heatmap
family_colors <- setNames(
```

```
    palette.colors(length(unique(ge$family)), "Okabe-Ito"),
    unique(ge$family)
)
Heatmap(
    pmin(assay(ge_signal, "counts")[, species_order], 5),
    name = "Copy number",
    col = c("white", "#fee0d2", "#fc9272", "#de2d26", "#a50f15", "#67000d"),
    cluster_columns = FALSE,
    row_split = factor(mod, levels = c(paste0("M", top_modules), "Other")),
    show_row_names = FALSE, show_row_dend = FALSE, cluster_row_slices = FALSE,
    row_title_rot = 0,
    top_annotation = HeatmapAnnotation(
        Family = ge$family[match(species_order, colnames(ge))],
        col = list(Family = family_colors)
    ),
    column_names_gp = gpar(fontsize = 9, fontface = "italic")
)
```

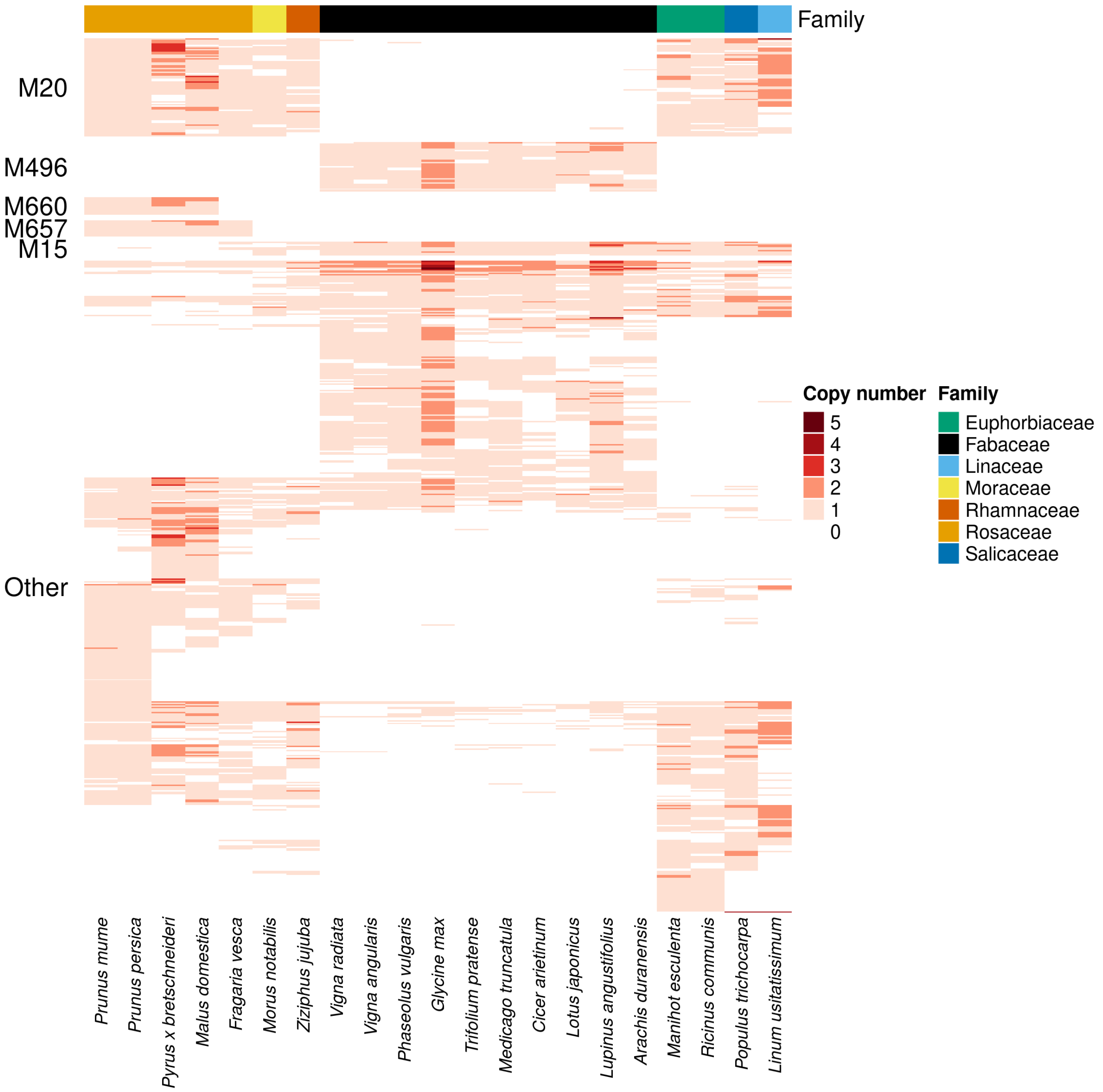

The heatmap shows that modules in the phylogenetic profile graph group synteny clusters with lineage-specific patterns of presence/absence (e.g., clusters that are specific to or lost in Fabaceae or Rosaceae). With all data in a single object, comparative genomics analyses that combine feature-level graphs (e.g., co-occurring gene families) and species-level graphs (e.g., phylogenies) become straightforward.

#### Session info

This document was created under the following conditions:

```
- Session info ---------------------------------------------------------------
 setting  value
 version  R version 4.5.2 (2025-10-31)
 os       Ubuntu 22.04.5 LTS
 system   x86_64, linux-gnu
 ui       X11
 language (EN)
 collate  en_US.UTF-8
 ctype    en_US.UTF-8
 tz       Europe/Brussels
 date     2026-10-24
 pandoc   3.6.3 @ /opt/quarto/bin/tools/ (via rmarkdown)
 quarto   1.8.25 @ /usr/local/bin/quarto

- Packages -------------------------------------------------------------------
 package              * version date (UTC) lib source
 ape                  * 5.8-1   2024-12-16 [1] CRAN (R 4.5.2)
 Biobase              * 2.70.0  2025-10-29 [1] Bioconductor 3.22 (R 4.5.2)
 BiocGenerics         * 0.56.0  2025-10-29 [1] Bioconductor 3.22 (R 4.5.2)
 ComplexHeatmap       * 2.26.1  2026-02-03 [1] Bioconductor 3.22 (R 4.5.2)
 dplyr                * 1.1.4   2023-11-17 [1] CRAN (R 4.5.2)
 forcats              * 1.0.1   2025-09-25 [1] CRAN (R 4.5.2)
 generics             * 0.1.4   2025-05-09 [1] CRAN (R 4.5.2)
 GenomicRanges        * 1.62.1  2025-12-08 [1] Bioconductor 3.22 (R 4.5.2)
 ggplot2              * 4.0.1   2025-11-14 [1] CRAN (R 4.5.2)
 ggraph               * 2.2.2   2025-08-24 [1] CRAN (R 4.5.2)
 GraphExperiment      * 1.1.2   2026-09-08 [1] Bioconductor
 here                 * 1.0.2   2025-09-15 [1] CRAN (R 4.5.2)
 igraph               * 2.2.1   2025-10-27 [1] CRAN (R 4.5.2)
 IRanges              * 2.44.0  2025-10-29 [1] Bioconductor 3.22 (R 4.5.2)
 lubridate            * 1.9.4   2024-12-08 [1] CRAN (R 4.5.2)
 MatrixGenerics       * 1.22.0  2025-10-29 [1] Bioconductor 3.22 (R 4.5.2)
 matrixStats          * 1.5.0   2025-01-07 [1] CRAN (R 4.5.2)
 purrr                * 1.2.1   2026-01-09 [1] CRAN (R 4.5.2)
 readr                * 2.1.6   2025-11-14 [1] CRAN (R 4.5.2)
 S4Vectors            * 0.48.0  2025-10-29 [1] Bioconductor 3.22 (R 4.5.2)
 Seqinfo              * 1.0.0   2025-10-29 [1] Bioconductor 3.22 (R 4.5.2)
 SingleCellExperiment * 1.32.0  2025-10-29 [1] Bioconductor 3.22 (R 4.5.2)
 stringr              * 1.6.0   2025-11-04 [1] CRAN (R 4.5.2)
 SummarizedExperiment * 1.40.0  2025-10-29 [1] Bioconductor 3.22 (R 4.5.2)
 syntenet             * 1.12.0  2025-10-30 [1] Bioconductor 3.22 (R 4.5.2)
 tibble               * 3.3.1   2026-01-11 [1] CRAN (R 4.5.2)
 tidyr                * 1.3.2   2025-12-19 [1] CRAN (R 4.5.2)
 tidyverse            * 2.0.0   2023-02-22 [1] CRAN (R 4.5.2)

 [1] /home/faalm/R/x86_64-pc-linux-gnu-library/4.5
 [2] /usr/local/lib/R/site-library
 [3] /usr/lib/R/site-library
 [4] /usr/lib/R/library
 * -- Packages attached to the search path.
```

### Use case 7: comparative methods on phylogenetic networks

Phylogenetic comparative methods (PCMs) typically assume that species relationships can be represented by a tree. However, reticulate evolutionary processes such as hybridization and introgression are common across the tree of life, and species relationships are often better described by phylogenetic networks, i.e., directed acyclic graphs (DAGs) in which some nodes (hybrid nodes) have more than one parent (Solís-Lemus and Ané 2016). Ignoring reticulation can bias estimates of evolutionary rates and lead to incorrect model selection (Morley, Lehmberg, and Kong 2026), but PCMs that explicitly model trait evolution on networks are still scarce, with notable exceptions such as the extension of Brownian motion (BM) models and phylogenetic regression to networks by Bastide et al. (2018).

Here, we will demonstrate how **GraphExperiment** can be used as the core data structure for comparative analyses using graphs (instead of trees) to represent species relationships. We will use data on *Xiphophorus* fishes (swordtails and platyfishes) from Bastide et al. (2018), which include phylogenetic networks inferred with SNaQ and two traits (Cui et al. 2013): the sword index (length of the sword, an extension of the caudal fin in males, relative to body size) and female preference for a sword.

```
library(here)
library(GraphExperiment)
library(tidyverse)
library(ggraph)
library(ComplexHeatmap)
library(patchwork)

# Load helper functions
source(here("code", "phylo_graph_utils.R"))

set.seed(123) # for reproducibility
```

The helper functions used in this chapter (`read_network_dag()`, `network_vcv()`, `vcv2graph()`, `graph2vcv()`, and `graph_gls()`) are defined in the file `code/phylo_graph_utils.R`, and their source code is available in the Appendices.

#### Data description

The data comprise a file with three time-calibrated networks (with 0, 1, and 3 reticulations) in extended Newick format, and a CSV file with trait data.

```
# Read networks
networks <- readLines(here("data", "xiphophorus_networks_calibrated.tre"))

# Read traits
traits <- read_csv(here("data", "xiphophorus_morphology_Cui_etal_2013.csv"))
head(traits)
```

```
# A tibble: 6 x 3
  tipnames    sword_index preference
  <chr>             <dbl>      <dbl>
1 Xalvarezi          0.65         NA
```

```
2 Xandersi          0.35       NA
3 Xbirchmanni       0.275      -0.33
4 Xclemenciae       0.564       0.44
5 Xcontinens        0.3        NA
6 Xcortezi          0.37       NA
```

We will represent species relationships as DAGs using `igraph` objects. Edges have attributes `length` (branch lengths) and `gamma` (inheritance probabilities, i.e., the proportion of the genome inherited from each parent; for tree edges, `gamma` is 1). The function `read_network_dag()` reads networks in extended Newick format and returns such DAGs. Here, we will use the network with 0 reticulations (i.e., a tree) and the one with 3 reticulations.

```
# Read tree (0 reticulations) and network (3 reticulations) as DAGs
dag_tree <- read_network_dag(networks[1])
dag_net <- read_network_dag(networks[3])

dag_net
```

```
IGRAPH be92156 DN-- 51 53 --
+ attr: name (v/c), is_tip (v/l), length (e/n), gamma (e/n)
+ edges from be92156 (vertex names):
 [1] node1 ->node2       node2 ->node3       node3 ->node4
 [4] node4 ->node5       node5 ->node6       node6 ->node7
 [7] node7 ->node8       node8 ->node9       node9 ->node10
[10] node10->Xgordoni    node10->Xmeyeri     node9 ->Xcouchianus
[13] node8 ->Xvariatus   node7 ->Xevelynae   node6 ->node11
[16] node11->Xxiphidium  node5 ->Xmilleri    node4 ->Xandersi
[19] node3 ->Xmaculatus  node2 ->node12      node12->node13
[22] node13->node14      node14->node15      node15->Xmontezumae
+ ... omitted several edges
```

```
# Hybrid nodes are nodes with in-degree > 1
V(dag_net)[degree(dag_net, mode = "in") > 1]
```

```
+ 3/51 vertices, named, from be92156:
[1] node12 node17 node27
```

```
# Hybrid edges (i.e., edges with gamma < 1)
igraph::as_data_frame(dag_net) |> filter(gamma < 1)
```

```
    from     to    length     gamma
1  node2 node12 10.221602 0.8346089
2 node16 node17  0.000000 0.8727956
3 node26 node27  1.633201 0.6307015
4 node11 node12  0.000000 0.1653911
5 node13 node17  1.735454 0.1272044
6 node24 node27  1.633200 0.3692985
```

Let's visualize the network. For that, we will compute node coordinates from the major tree (i.e., the tree obtained by keeping only the hybrid edges with the largest inheritance probabilities), with node positions on the x-axis representing time since the root. Then, we will add minor hybrid edges on top.

```
# Get major tree by removing minor hybrid edges
dag_major <- delete_edges(dag_net, E(dag_net)[gamma < 0.5])
```

```r
root <- V(dag_major)[degree(dag_major, mode = "in") == 0]

# Horizontal positions: distance from the root
depth <- distances(dag_major, v = root, mode = "out", weights =
 ↪  E(dag_major)$length)[1, ]

# Vertical positions: tips ordered by depth-first search, and internal nodes
# in the middle of their children (visiting children before parents)
dfs_order <- names(dfs(dag_major, root, mode = "out")$order)
pos <- setNames(rep(NA, vcount(dag_major)), V(dag_major)$name)
pos[V(dag_major)$is_tip] <- rank(match(V(dag_major)$name[V(dag_major)$is_tip],
 ↪  dfs_order))
pos <- Reduce(function(pos, v) {
    if(is.na(pos[v])) pos[v] <- mean(pos[neighbors(dag_major, v, mode =
     ↪  "out")$name])
    pos
}, rev(dfs_order), init = pos)

# Plot network with the coordinates of the major tree
ggraph(dag_net, layout = "manual", x = depth[V(dag_net)$name], y =
 ↪  pos[V(dag_net)$name]) +
    geom_edge_diagonal(aes(filter = gamma == 1), color = "grey50", flipped = TRUE) +
    geom_edge_link(
        aes(filter = gamma < 1, color = gamma),
        arrow = arrow(length = unit(2, "mm"), type = "closed"),
        end_cap = circle(1, "mm"), linewidth = 0.8
    ) +
    geom_node_text(
        aes(filter = is_tip, label = name), hjust = -0.1,
        size = 3, fontface = "italic"
    ) +
    scale_edge_color_gradient(low = "#fcbba1", high = "firebrick") +
    scale_x_continuous(expand = expansion(mult = c(0.02, 0.2))) +
    labs(edge_color = "Inheritance\nprobability (hybrid edges)") +
    theme_graph(base_family = "sans")
```

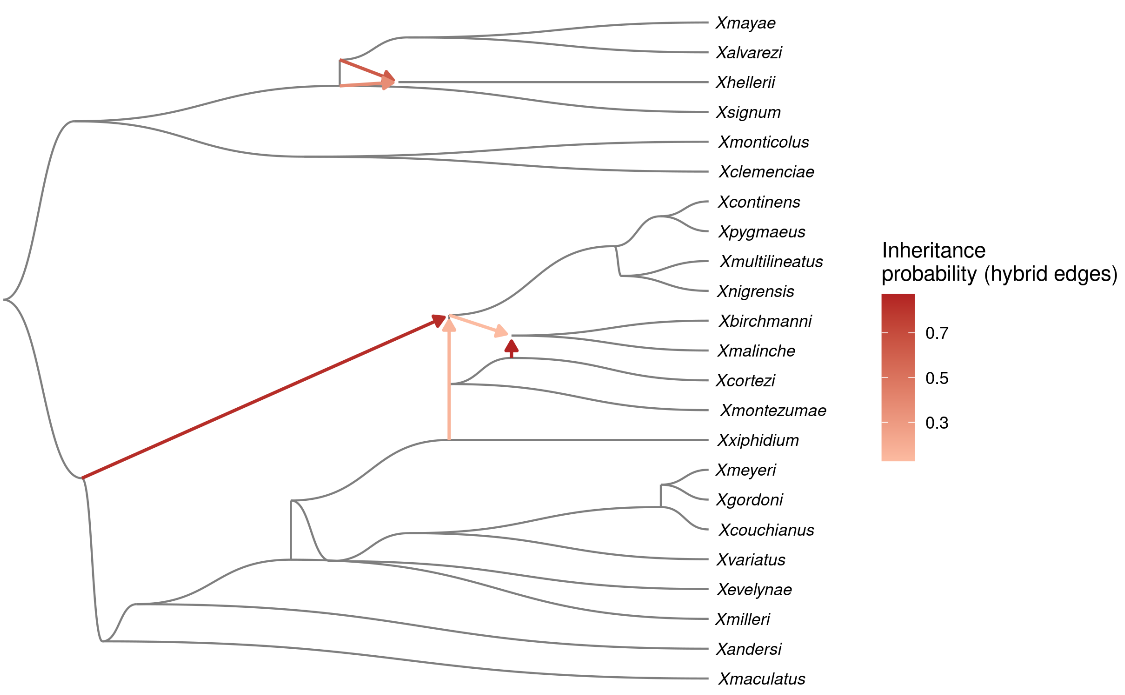

#### Storing species graphs in a `GraphExperiment` object

The trait data will be stored in a matrix with traits in rows and species in columns. One species (*X. nezahualcoyotl*) is not in the network, so we will remove it. We will also add a column with clade information (northern swordtails, southern swordtails, and platyfishes) to `colData`.

```
# Get trait matrix
tips <- V(dag_net)$name[V(dag_net)$is_tip]
trait_mat <- traits |>
    filter(tipnames %in% tips) |>
    column_to_rownames("tipnames") |>
    as.matrix() |>
    t()

trait_mat[, 1:5]
```

```
            Xalvarezi Xandersi Xbirchmanni Xclemenciae Xcontinens
sword_index      0.65     0.35       0.275       0.564        0.3
preference         NA       NA      -0.330       0.440         NA
```

```
# Create species metadata
northern <- c(
    "Xbirchmanni", "Xcontinens", "Xcortezi", "Xmalinche", "Xmontezumae",
    "Xmultilineatus", "Xnigrensis", "Xpygmaeus"
)
southern <- c(
    "Xalvarezi", "Xclemenciae", "Xhellerii", "Xmayae", "Xmonticolus", "Xsignum"
)
species_md <- data.frame(
```

```r
    row.names = colnames(trait_mat),
    clade = case_when(
        colnames(trait_mat) %in% northern ~ "Northern swordtails",
        colnames(trait_mat) %in% southern ~ "Southern swordtails",
        TRUE ~ "Platyfishes"
    )
)
```

Nodes of graphs in `colGraphs` must match column names (here, species) *exactly*. Since our DAGs contain internal nodes (ancestors) in addition to tips (extant species), they cannot be stored in the `colGraphs` slot as they are. Instead, we will store in `colGraphs` the *species-species* graphs that comparative methods actually use: the variance-covariance structure expected under BM. In these graphs, nodes are extant species, edge weights are the covariances between species (i.e., their shared evolutionary history), and a node attribute (`variance`) stores the variances (i.e., root-to-tip distances). These are computed with `network_vcv()`, which implements the recursive algorithm of Bastide et al. (2018) on the DAG: nodes are visited in topological order, and the trait of each hybrid node is the average of its parents' traits weighted by inheritance probabilities.

> **Note**
>
> Native support for graphs containing nodes that are not observations (e.g., internal nodes of phylogenies and phylogenetic networks) is planned for future versions of **GraphExperiment**. For now, full DAGs can be stored in the `metadata` slot, as we will do below.

Instead, we will store in `colGraphs` the *species-species* graphs that comparative methods actually use: the variance-covariance structure expected under BM. In these graphs, nodes are extant species, edge weights are the covariances between species (i.e., their shared evolutionary history), and a node attribute (`variance`) stores the variances (i.e., root-to-tip distances). These are computed with `network_vcv()`, which implements the recursive algorithm of Bastide et al. (2018) on the DAG: nodes are visited in topological order, and the trait of each hybrid node is the average of its parents' traits weighted by inheritance probabilities.

```r
# Get variance-covariance matrices under BM
vcv_tree <- network_vcv(dag_tree)
vcv_net <- network_vcv(dag_net)

# Sanity check: for the tree, we get the same as `ape::vcv()`
vcv_ape <- ape::vcv(ape::read.tree(text = networks[1]))
all.equal(vcv_tree, vcv_ape[rownames(vcv_tree), colnames(vcv_tree)])
```

```
[1] TRUE
```

```r
# Create a `GraphExperiment` object
species <- colnames(trait_mat)
ge <- GraphExperiment(
    assays = list(traits = trait_mat),
    colData = species_md,
    colGraphs = list(
        tree_vcv = vcv2graph(vcv_tree[species, species]),
        network_vcv = vcv2graph(vcv_net[species, species])
    ),
    metadata = list(dag_tree = dag_tree, dag_net = dag_net)
)

ge
```

```
class: GraphExperiment
dim: 2 23
metadata(2): dag_tree dag_net
```

```
assays(1): traits
rownames(2): sword_index preference
rowData names(0):
colnames(23): Xalvarezi Xandersi ... Xvariatus Xxiphidium
colData names(3): clade tree_vcv__variance network_vcv__variance
reducedDimNames(0):
mainExpName: NULL
altExpNames(0):
rowGraphs(0):
colGraphs(2): tree_vcv network_vcv
```

As node attributes are automatically added to `colData`, we can easily see that, under a BM model on the network, hybrid species have smaller variances than expected under a tree. This is because the trait of a hybrid is a weighted average of its parents' traits.

```
colData(ge) |>
    as.data.frame() |>
    mutate(diff = network_vcv__variance - tree_vcv__variance) |>
    arrange(diff) |>
    head()
```

```
                               clade tree_vcv__variance network_vcv__variance
Xbirchmanni    Northern swordtails           19.63128              16.42402
Xmalinche      Northern swordtails           19.63128              16.42402
Xcortezi       Northern swordtails           19.63128              16.80936
Xmontezumae    Northern swordtails           19.63128              16.80936
Xmultilineatus Northern swordtails           19.63128              16.80937
Xnigrensis     Northern swordtails           19.63128              16.80937
                    diff
Xbirchmanni    -3.207260
Xmalinche      -3.207260
Xcortezi       -2.821913
Xmontezumae    -2.821913
Xmultilineatus -2.821912
Xnigrensis     -2.821912
```

Let's visualize the difference in shared evolutionary history between species under the network and under the tree.

```
# Get difference between covariance matrices from `colGraphs`
cov_diff <- graph2vcv(colGraph(ge, "network_vcv")) -
    graph2vcv(colGraph(ge, "tree_vcv"))

# Plot heatmap, with species annotated by clade (from `colData`)
clade_colors <- c(
    "Northern swordtails" = "#1b9e77", "Southern swordtails" = "#d95f02",
    "Platyfishes" = "#7570b3"
)
ha <- HeatmapAnnotation(
    Clade = ge$clade, col = list(Clade = clade_colors)
)
Heatmap(
    cov_diff, name = "Covariance\n(network - tree)",
    top_annotation = ha,
    col = circlize::colorRamp2(c(-4, 0, 4), c("steelblue", "white", "firebrick")),
    row_names_gp = gpar(fontsize = 8, fontface = "italic"),
    column_names_gp = gpar(fontsize = 8, fontface = "italic")
)
```

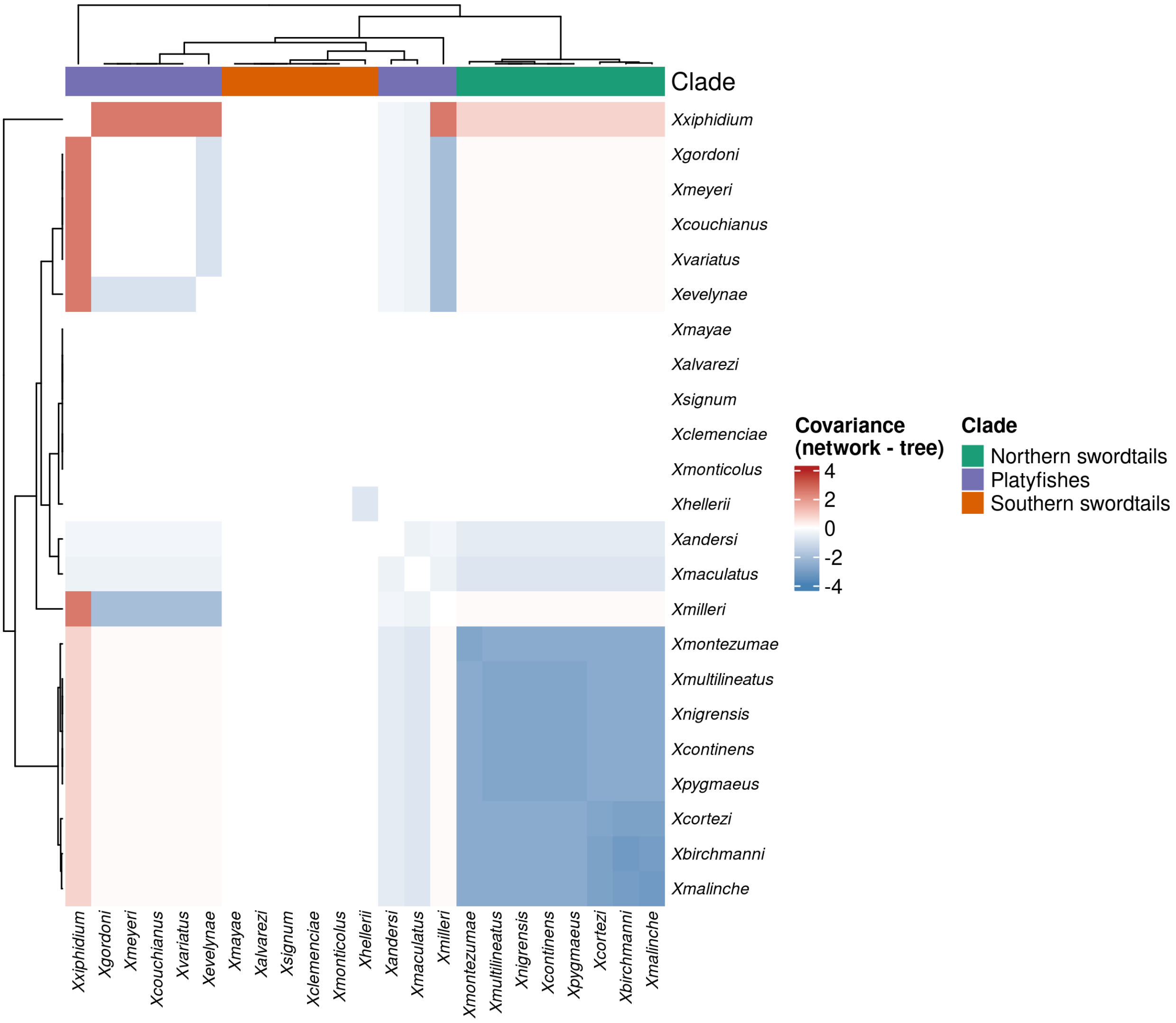

The heatmap shows that, by accounting for the hybrid origin of northern swordtails (with genetic contribution from the lineage leading to *X. xiphidium*; see network plot above), covariances between northern swordtails and platyfishes increase, while covariances among northern swordtails decrease.

#### Comparative analyses using graphs

Now, we will fit models of trait evolution using the species graphs stored in `colGraphs`. The function `graph_gls()` takes a `GraphExperiment` object, the name of a trait (row), and the name of a graph in `colGraphs`, and fits a phylogenetic generalized least squares (GLS) model under BM (or under a Pagel's $\lambda$ model, in which covariances are multiplied by $\lambda$ to measure phylogenetic signal). Importantly, `graph_gls()` does not care whether the graph was derived from a tree, a network, or any other source (e.g., gene-sharing networks, ancestral recombination graphs, etc.); it only needs a species-species graph.

```
# Fit models for each combination of trait, graph, and model
fits <- expand_grid(
    trait = rownames(ge),
    graph = colGraphNames(ge),
    model = c("BM", "lambda")
) |>
    pmap(\(trait, graph, model) graph_gls(ge, trait, graph, model)) |>
    bind_rows()
```

```r
fits |> mutate(across(where(is.numeric), \(x) signif(x, 3)))
```

```
        trait       graph  model  n root_state  sigma2 lambda  logLik    AIC
1 sword_index    tree_vcv     BM 23      0.461 0.00266      1  9.2100 -14.40
2 sword_index    tree_vcv lambda 23      0.461 0.00266      1  9.2100 -12.40
3 sword_index network_vcv     BM 23      0.464 0.00275      1  8.9700 -13.90
4 sword_index network_vcv lambda 23      0.464 0.00275      1  8.9700 -11.90
5  preference    tree_vcv     BM 10      0.347 0.00508      1 -0.0156   4.03
6  preference    tree_vcv lambda 10      0.347 0.00508      1 -0.0160   6.03
7  preference network_vcv     BM 10      0.320 0.00566      1 -0.3630   4.73
8  preference network_vcv lambda 10      0.320 0.00566      1 -0.3630   6.73
```

Both traits show strong phylogenetic signal ($\lambda$ estimates at the upper bound), in line with Bastide et al. (2018). For these two traits, models based on the tree and on the network fit the data similarly well (differences in AIC $< 2$), indicating that the inferred reticulations do not change conclusions about their mode of evolution. Comparisons like this are straightforward when all candidate species graphs are stored in the same object. Finally, let's store the best model for each trait in `rowData`, so that results stay synchronized with the data.

```r
# Get best model for each trait and add to `rowData`
best <- fits |>
    group_by(trait) |>
    slice_min(AIC, n = 1) |>
    ungroup()

rowData(ge)$best_graph <- best$graph[match(rownames(ge), best$trait)]
rowData(ge)$best_model <- best$model[match(rownames(ge), best$trait)]
rowData(ge)$lambda <- best$lambda[match(rownames(ge), best$trait)]

rowData(ge)
```

```
DataFrame with 2 rows and 3 columns
             best_graph  best_model    lambda
            <character> <character> <numeric>
sword_index    tree_vcv          BM         1
preference     tree_vcv          BM         1
```

In summary, by storing traits, species metadata, and species graphs in a single object, **GraphExperiment** provides a standard data structure for developing new comparative methods based on graphs. Developers only need to write functions that take a `GraphExperiment` object and the name of a graph in `colGraphs` as input, as we did with `graph_gls()`, and users can easily compare results obtained with different representations of species relationships (e.g., trees, networks, gene-sharing networks, or any other species graph).

#### Session info

This document was created under the following conditions:

```
- Session info ---------------------------------------------------------------
 setting  value
 version  R version 4.5.2 (2025-10-31)
 os       Ubuntu 22.04.5 LTS
 system   x86_64, linux-gnu
 ui       X11
 language (EN)
 collate  en_US.UTF-8
 ctype    en_US.UTF-8
```

```
 tz       Europe/Brussels
 date     2026-10-24
 pandoc   3.6.3 @ /opt/quarto/bin/tools/ (via rmarkdown)
 quarto   1.8.25 @ /usr/local/bin/quarto

- Packages ---------------------------------------------------------------------
 package              * version date (UTC) lib source
 Biobase              * 2.70.0  2025-10-29 [1] Bioconductor 3.22 (R 4.5.2)
 BiocGenerics         * 0.56.0  2025-10-29 [1] Bioconductor 3.22 (R 4.5.2)
 ComplexHeatmap       * 2.26.1  2026-02-03 [1] Bioconductor 3.22 (R 4.5.2)
 dplyr                * 1.1.4   2023-11-17 [1] CRAN (R 4.5.2)
 forcats              * 1.0.1   2025-09-25 [1] CRAN (R 4.5.2)
 generics             * 0.1.4   2025-05-09 [1] CRAN (R 4.5.2)
 GenomicRanges        * 1.62.1  2025-12-08 [1] Bioconductor 3.22 (R 4.5.2)
 ggplot2              * 4.0.1   2025-11-14 [1] CRAN (R 4.5.2)
 ggraph               * 2.2.2   2025-08-24 [1] CRAN (R 4.5.2)
 GraphExperiment      * 1.1.2   2026-09-08 [1] Bioconductor
 here                 * 1.0.2   2025-09-15 [1] CRAN (R 4.5.2)
 igraph               * 2.2.1   2025-10-27 [1] CRAN (R 4.5.2)
 IRanges              * 2.44.0  2025-10-29 [1] Bioconductor 3.22 (R 4.5.2)
 lubridate            * 1.9.4   2024-12-08 [1] CRAN (R 4.5.2)
 MatrixGenerics       * 1.22.0  2025-10-29 [1] Bioconductor 3.22 (R 4.5.2)
 matrixStats          * 1.5.0   2025-01-07 [1] CRAN (R 4.5.2)
 patchwork            * 1.3.2   2025-08-25 [1] CRAN (R 4.5.2)
 purrr                * 1.2.1   2026-01-09 [1] CRAN (R 4.5.2)
 readr                * 2.1.6   2025-11-14 [1] CRAN (R 4.5.2)
 S4Vectors            * 0.48.0  2025-10-29 [1] Bioconductor 3.22 (R 4.5.2)
 Seqinfo              * 1.0.0   2025-10-29 [1] Bioconductor 3.22 (R 4.5.2)
 SingleCellExperiment * 1.32.0  2025-10-29 [1] Bioconductor 3.22 (R 4.5.2)
 stringr              * 1.6.0   2025-11-04 [1] CRAN (R 4.5.2)
 SummarizedExperiment * 1.40.0  2025-10-29 [1] Bioconductor 3.22 (R 4.5.2)
 tibble               * 3.3.1   2026-01-11 [1] CRAN (R 4.5.2)
 tidyr                * 1.3.2   2025-12-19 [1] CRAN (R 4.5.2)
 tidyverse            * 2.0.0   2023-02-22 [1] CRAN (R 4.5.2)

 [1] /home/faalm/R/x86_64-pc-linux-gnu-library/4.5
 [2] /usr/local/lib/R/site-library
 [3] /usr/lib/R/site-library
 [4] /usr/lib/R/library
 * -- Packages attached to the search path.

--------------------------------------------------------------------------------
```

Vanderaa, Christophe, and Laurent Gatto. 2023a. "The Current State of Single-cell Proteomics Data Analysis."

*Current Protocols* 3 (1). https://doi.org/10.1002/cpz1.658.

———. 2023b. "Revisiting the Thorny Issue of Missing Values in Single-Cell Proteomics." *Journal of Proteome Research* 22 (9): 2775–84. https://doi.org/10.1021/acs.jproteome.3c00227.

———. 2025. "Scplainer: Using Linear Models to Understand Mass Spectrometry-Based Single-Cell Proteomics Data." *Genome Biology* 26 (1). https://doi.org/10.1186/s13059-025-03713-4.

Wright, Kevin. 2026. *Nipals: Principal Components Analysis Using NIPALS or Weighted EMPCA, with Gram-Schmidt Orthogonalization*. https://doi.org/10.32614/CRAN.package.nipals.

Xu, Chen, and Zhengchang Su. 2015. "Identification of Cell Types from Single-Cell Transcriptomes Using a Novel Clustering Method." *Bioinformatics* 31 (12): 1974–80. https://doi.org/10.1093/bioinformatics/btv088.

Zeisel, Amit, Ana B. Muñoz-Manchado, Simone Codeluppi, Peter Lönnerberg, Gioele La Manno, Anna Juréus, Sueli Marques, et al. 2015. "Cell Types in the Mouse Cortex and Hippocampus Revealed by Single-Cell RNA-Seq." *Science* 347 (6226): 1138–42. https://doi.org/10.1126/science.aaa1934.

Zhao, Tao, and M. Eric Schranz. 2019. "Network-Based Microsynteny Analysis Identifies Major Differences and Genomic Outliers in Mammalian and Angiosperm Genomes." *Proceedings of the National Academy of Sciences* 116 (6): 2165–74. https://doi.org/10.1073/pnas.1801757116.

Zhao, Tao, Arthur Zwaenepoel, Jia-Yu Xue, Shu-Min Kao, Zhen Li, M. Eric Schranz, and Yves Van de Peer. 2021. "Whole-Genome Microsynteny-Based Phylogeny of Angiosperms." *Nature Communications* 12 (1). https://doi.org/10.1038/s41467-021-23665-0.

### Appendices

Here, you can find other pieces of code (Bash and R scripts) that were used in the main chapters. These pieces of code were stored in separate files to make the main text (i.e., book chapters) more readable.

#### Helper functions for phylogenetic networks

Below is the source code of the helper functions used in Use case 7 to read phylogenetic networks as `igraph` DAGs, compute Brownian motion variance-covariance matrices on networks, convert them to and from species graphs, and fit phylogenetic GLS models using graphs stored in the `colGraphs` slot of a `GraphExperiment` object. These functions are stored in the file `code/phylo_graph_utils.R`.

```r
#' Read a phylogenetic network in extended Newick format as an igraph DAG
#'
#' @param text Character with a network in extended Newick format,
#' with inheritance probabilities (gamma) after a double colon
#' (e.g., "#H1:0.5::0.3"), as written by SNaQ/PhyloNetworks.
#'
#' @return A directed `igraph` object (a DAG) with edge attributes
#' 'length' (branch length) and 'gamma' (inheritance probability), and
#' vertex attribute 'is_tip'.
#'
read_network_dag <- function(text) {

    if(grepl("#H", text)) {
        net <- ape::read.evonet(text = text)
    } else { # tree (no reticulations)
        net <- ape::read.tree(text = text)
        net$reticulation <- matrix(integer(0), ncol = 2)
    }
    ntip <- length(net$tip.label)
    node_names <- c(net$tip.label, paste0("node", seq_len(net$Nnode)))

    # Tree edges (including major hybrid edges)
    edges <- data.frame(
        from = node_names[net$edge[, 1]],
        to = node_names[net$edge[, 2]],
        length = net$edge.length,
        gamma = 1
    )

    # Get length and gamma of hybrid edges from the Newick string
    hyb <- regmatches(text, gregexpr("\\)?#H[0-9]+:[^,);]*", text))[[1]]
    hyb_df <- data.frame(
        label = sub(":.*", "", sub("^\\)", "", hyb)),
        defining = startsWith(hyb, ")"), # occurrence with a subtree
        length = as.numeric(sapply(strsplit(hyb, ":"), `[`, 2)),
        gamma = as.numeric(sapply(strsplit(hyb, "::"), `[`, 2))
    )

    # Update major hybrid edges (parent edge of each hybrid node)
    hnode_labels <- net$node.label[net$reticulation[, 2] - ntip]
```

```r
    hnodes <- node_names[net$reticulation[, 2]]
    major_gamma <- vapply(hnode_labels, function(h) {
        hyb_df$gamma[hyb_df$label == h & hyb_df$defining]
    }, numeric(1))
    idx <- match(edges$to, hnodes)
    edges$gamma[!is.na(idx)] <- major_gamma[idx[!is.na(idx)]]

    # Add minor hybrid edges
    minor_edges <- lapply(seq_len(nrow(net$reticulation)), function(i) {
        minor <- hyb_df[hyb_df$label == hnode_labels[i] & !hyb_df$defining, ]
        data.frame(
            from = node_names[net$reticulation[i, 1]],
            to = node_names[net$reticulation[i, 2]],
            length = minor$length,
            gamma = minor$gamma
        )
    })
    edges <- rbind(edges, do.call(rbind, minor_edges))

    # Create DAG
    dag <- igraph::graph_from_data_frame(
        edges, directed = TRUE,
        vertices = data.frame(
            name = node_names, is_tip = seq_along(node_names) <= ntip
        )
    )

    return(dag)
}

#' Compute the Brownian motion variance-covariance matrix on a phylogenetic DAG
#'
#' Implements the recursive algorithm of Bastide et al. (2018, Syst Biol):
#' nodes are visited in topological order (parents before children); for tree
#' nodes, covariances are inherited from the parent; for hybrid nodes,
#' the trait is a weighted average (weights = gamma) of the parents' traits.
#' Works for trees (DAGs without hybrid nodes) and networks.
#'
#' @param dag A directed `igraph` object as returned by `read_network_dag()`.
#'
#' @return A tips x tips variance-covariance matrix.
#'
network_vcv <- function(dag) {

    ord <- igraph::as_ids(igraph::topo_sort(dag, mode = "out"))
    n <- igraph::vcount(dag)
    nodes <- igraph::V(dag)$name
    vcv <- matrix(0, n, n, dimnames = list(nodes, nodes))

    # Update covariances of node `v` given those of previously visited nodes
    update_vcv <- function(vcv, i) {
        v <- ord[i]
        visited <- ord[seq_len(i - 1)]
        in_edges <- igraph::incident(dag, v, mode = "in")
        if(length(in_edges) == 0) return(vcv) # root

        parents <- igraph::tail_of(dag, in_edges)$name
        g <- in_edges$gamma
```

```r
        l <- in_edges$length

        # Covariance with previously visited nodes
        vcv[v, visited] <- vcv[visited, v] <- colSums(
            g * vcv[parents, visited, drop = FALSE]
        )

        # Variance
        vcv[v, v] <- sum(g^2 * (diag(vcv)[parents] + l)) +
            sum(outer(g, g) * vcv[parents, parents] * (1 - diag(length(g))))

        return(vcv)
    }
    vcv <- Reduce(update_vcv, seq_along(ord), init = vcv)

    tips <- nodes[igraph::V(dag)$is_tip]
    return(vcv[tips, tips])
}

#' Convert a variance-covariance matrix to a weighted species graph
#'
#' @param vcv A species x species variance-covariance matrix.
#'
#' @return An undirected `igraph` object with edge weights representing
#' covariances (i.e., shared evolutionary history), and vertex attribute
#' 'variance' storing the diagonal of `vcv` (i.e., root-to-tip distances).
#'
vcv2graph <- function(vcv) {
    g <- igraph::graph_from_adjacency_matrix(
        vcv, mode = "undirected", weighted = TRUE, diag = FALSE
    )
    igraph::V(g)$variance <- diag(vcv)[igraph::V(g)$name]
    return(g)
}

#' Extract a variance-covariance matrix from a species graph
#'
#' @param graph An undirected `igraph` object as returned by `vcv2graph()`.
#'
#' @return A species x species variance-covariance matrix.
#'
graph2vcv <- function(graph) {
    C <- as.matrix(igraph::as_adjacency_matrix(graph, attr = "weight"))
    diag(C) <- igraph::V(graph)$variance
    return(C)
}

#' Fit a phylogenetic GLS model (intercept only) using a species graph
#'
#' Models a trait as evolving under Brownian motion (BM), with the
#' covariance among species obtained from a `colGraph` of a
#' `GraphExperiment` object. Optionally, Pagel's lambda is estimated
#' by maximum likelihood.
#'
#' @param ge A `GraphExperiment` object with traits in rows and
#' species in columns.
```

```r
#' @param trait Character, name of the trait (row) to model.
#' @param graph Character, name of the graph in `colGraphs` with the
#' covariance structure (as returned by `vcv2graph()`).
#' @param model Character, either "BM" or "lambda".
#' @param assay Character, name of the assay to use.
#'
#' @return A data frame with the estimated parameters, log-likelihood, and AIC.
#'
graph_gls <- function(ge, trait, graph, model = "BM", assay = 1) {

    vals <- SummarizedExperiment::assay(ge, assay)[trait, ]
    vals <- vals[!is.na(vals)]
    C <- graph2vcv(GraphExperiment::colGraph(ge, graph))[names(vals), names(vals)]
    n <- length(vals)
    dm <- matrix(1, n, 1) # design matrix (intercept only)

    # Profile log-likelihood for a given lambda
    loglik <- function(lambda) {
        Cl <- C * lambda
        diag(Cl) <- diag(C)
        Ci <- solve(Cl)
        beta <- solve(t(dm) %*% Ci %*% dm, t(dm) %*% Ci %*% vals)
        r <- vals - dm %*% beta
        sigma2 <- as.numeric(t(r) %*% Ci %*% r) / n
        ll <- -n / 2 * log(2 * pi * sigma2) -
            as.numeric(determinant(Cl)$modulus) / 2 - n / 2
        list(ll = ll, beta = as.numeric(beta), sigma2 = sigma2)
    }

    lambda <- 1
    if(model == "lambda") {
        lambda <- optimize(function(l) loglik(l)$ll, c(0, 1), maximum =
 ↪  TRUE)$maximum
    }
    fit <- loglik(lambda)
    k <- 2 + (model == "lambda") # number of parameters

    res <- data.frame(
        trait = trait, graph = graph, model = model, n = n,
        root_state = fit$beta, sigma2 = fit$sigma2, lambda = lambda,
        logLik = fit$ll, AIC = -2 * fit$ll + 2 * k
    )
    return(res)
}
```

#### Session info

This document was created under the following conditions:

```
- Session info ---------------------------------------------------------------
 setting  value
 version  R version 4.5.2 (2025-10-31)
 os       Ubuntu 22.04.5 LTS
 system   x86_64, linux-gnu
 ui       X11
 language (EN)
 collate  en_US.UTF-8
 ctype    en_US.UTF-8
```

```
 tz       Europe/Brussels
 date     2026-10-24
 pandoc   3.6.3 @ /opt/quarto/bin/tools/ (via rmarkdown)
 quarto   1.8.25 @ /usr/local/bin/quarto

──────────────────────────────────────────────────────────────────────────────
```
